# Genomics of Secondarily Temperate Adaptation in the Only Non-Antarctic Icefish

**DOI:** 10.1101/2022.08.13.503862

**Authors:** Angel G. Rivera-Colón, Niraj Rayamajhi, Bushra Fazal Minhas, Giovanni Madrigal, Kevin T. Bilyk, Veronica Yoon, Mathias Hüne, Susan Gregory, C.-H. Christina Cheng, Julian M. Catchen

**Affiliations:** Department of Evolution, Ecology, and Behavior, University of Illinois at Urbana-Champaign, Urbana, Illinois, USA; Informatics Program, University of Illinois at Urbana-Champaign, Urbana, Illinois, USA; Department of Biology, Montclair State University, Montclair, New Jersey, USA; Centro de Investigación para la Conservación de los Ecosistemas Australes, Punta Arenas, Chile; British Antarctic Survey, High Cross, Madingley Road, Cambridge, CB3 0ET, UK; Government of South Georgia and the South Sandwich Islands, Stanley FIQQ 1ZZ, Falklands

**Author notes:** Corresponding Author: Julian M. Catchen.

## Abstract

White-blooded Antarctic icefishes, a family within the adaptive radiation of Antarctic notothenioid fishes, are an example of extreme biological specialization to both the chronic cold of the Southern Ocean and life without hemoglobin. As a result, icefishes display derived physiology that limits them to the cold and highly oxygenated Antarctic waters. Against these constraints, remarkably one species, the pike icefish *Champsocephalus esox*, successfully colonized temperate South American waters. To study the genetic mechanisms underlying secondarily temperate adaptation in icefishes, we generated chromosome-level genome assemblies of both *C. esox* and its Antarctic sister species, *Champsocephalus gunnari*. The *C. esox* genome is similar in structure and organization to that of its Antarctic congener; however, we observe evidence of chromosomal rearrangements coinciding with regions of elevated genetic divergence in pike icefish populations. We also find several key biological pathways under selection, including genes related to mitochondria and vision, highlighting candidates behind temperate adaptation in *C. esox*. Substantial antifreeze glycoprotein (AFGP) pseudogenization has occurred in the pike icefish, likely due to relaxed selection following ancestral escape from Antarctica. The canonical *AFGP* locus organization is conserved in *C. esox* and *C. gunnari*, but both show a translocation of two *AFGP* copies to a separate locus, previously unobserved in cryonotothenioids. Altogether, the study of this secondarily temperate species provides an insight into the mechanisms underlying adaptation to ecologically disparate environments in this otherwise highly specialized group.

## Introduction

Antarctic notothenioid fish, or cryonotothenioids, have experienced extreme biological specialization, evolving and diversifying in the chronically cold environment of the Southern Ocean. Following Antarctica’s thermal and geographical isolation over the last 35 million years (Mya) (Eastman 2005), cryonotothenioids evolved molecular and physiological adaptations to the extreme cold, most notably antifreeze glycoproteins (AFGPs) (Chen et al. 1997; DeVries and Cheng 2005) that allowed them to thrive in an otherwise inhospitable environment. As other Antarctic fish taxa became extinct, this group displayed rapid rates of diversification and underwent an adaptive radiation (Matschiner et al. 2011; Near et al. 2012; Colombo et al. 2015) dominating modern fish fauna in the Southern Ocean and playing a key role in the ecosystem’s food web (La Mesa et al. 2004). However, cold specialization has its tradeoffs. Cryonotothenioids are stenothermal, with physiological performance limited to low and narrow temperature windows (Beers and Sidell 2011), and are unable to mount the classic heat shock response (Hofmann et al. 2000; Bilyk and Cheng 2014; Bilyk et al. 2018).

Among Antarctic notothenioids, further specialization is observed in the family of the white-blooded icefishes (Channichthyidae). Characterized by a complete loss of both hemoglobin and red blood cells (Sidell and O’Brien 2006), icefish have evolved various mechanisms to compensate for this lack of active oxygen transport including larger hearts and thus greater cardiac stroke volume (Sidell and O’Brien 2006), changes in the morphology and density of mitochondria (Johnston et al. 1998; O’Brien and Mueller 2010), a novel molecular method to increase CO_2_ excretion (Harter et al. 2018), and increased cellular lipids (Palmerini et al. 2009). Even after the evolution of these phenotypes, icefish still exhibit limited cardiac performance, with less than 10% of the oxygen carrying capacity of red-blooded notothenioids (Holeton 1970) and far larger cardiac energy expenditures (Tota et al. 1991; Tota and Gattuso 1996). Given these biological limitations, the loss of hemoglobin in icefish was often considered to be maladaptive (Sidell and O’Brien 2006), although recent work has proposed alternative hypotheses related to adaptive changes in the presence of sparse iron and alternative functional roles for partial hemoglobin fragments (Corliss et al. 2019). Regardless of the evolutionary mechanism behind the loss, the derived physiology stemming from the lack of hemoglobin is predicted to limit icefishes’ adaptive potential to migrate to warmer waters or to respond to rapidly changing ocean temperatures.

Despite these constraints, a single icefish lineage succeeded in establishing itself north of the Antarctic Polar Front (Stankovic et al. 2002; Kock 2005). The pike icefish, *Champsocephalus esox*, is distributed along the Patagonia coast and Strait of Magellan in South America, and around the Falkland Islands (Fig. 1), non-freezing and seasonally variable environments that are significantly warmer, and much lower in oxygen concentration than the Southern Ocean. *C. esox* shares the defining traits of the Antarctic icefish clade, including the lack of hemoglobin (Grove et al. 2004; Kock 2005), loss of myoglobin (like its Antarctic congener, *Champsocephalus gunnari* (Grove et al. 2004)), and with AFGP coding sequences apparently still present in its DNA (Miya et al. 2016). This prompts the question: how does adaptation and survival in a temperate environment occur from a highly derived and cold-specialized Antarctic ancestral state? The pike icefish presents an ideal model to address this evolutionary question. Its recent divergence (~1.6 Mya (Stankovic et al. 2002; Near et al. 2012; Dornburg et al. 2017)) from its Antarctic sister species, the mackerel icefish *C. gunnari*, enables a direct comparison of closely related Antarctic and temperate genomes. *C. gunnari* serves as a proxy of the Antarctic ancestral state, against which we can identify genetic changes in *C. esox* related to the adaptation to a new temperate environment.

**Figure 1.**
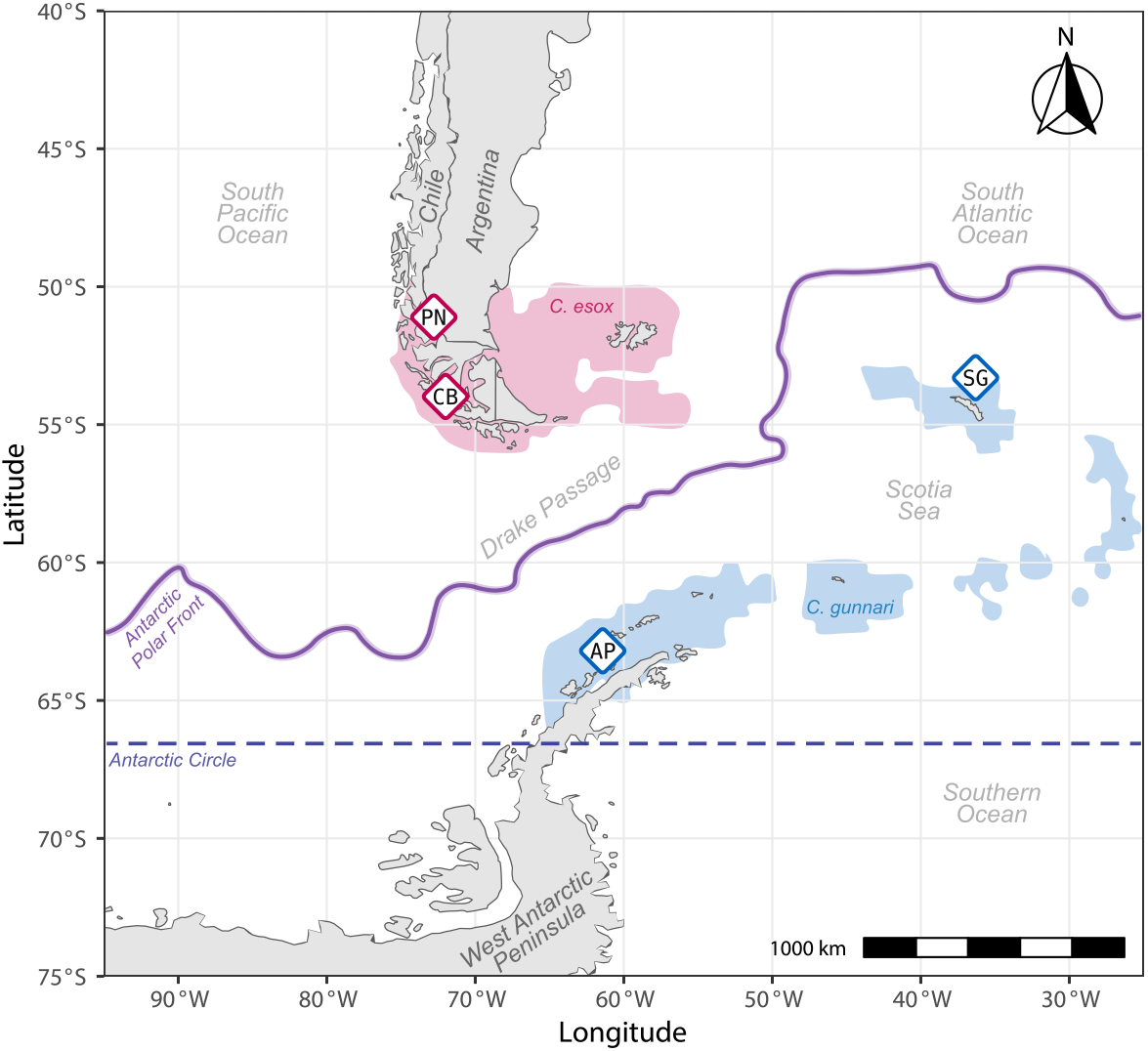
Species ranges and sampling of temperate and Antarctic icefish species. Map showing the approximate species range of the temperate pike icefish *C. esox* (in red) and the Antarctic mackerel icefish *C. gunnari* (in blue) (AquaMaps 2019). The approximate boundary of the Antarctic Polar Front is shown in the solid purple line (Orsi et al. 1995). *C. esox* individuals were sampled form Puerto Natales, Chile (PN, n=16) and Canal Bárbara, Chile (CB, n=12), while the *C. gunnari* individuals were collected from the Gerlache Strait in the West Antarctic Peninsula (AP, n=20) and South Georgia island (SG, n=48). Genomes for *C. esox* and *C. gunnari* were assembled from individuals collected from the PN and AP populations, respectively.

*C. esox* remains a poorly researched species. The few published studies provide some examples of phenotypic differences in *C. esox* compared to other icefish, including different reproductive biology (Calvo et al. 1999), increased mitochondrial densities (Johnston et al. 1998), and mitochondrial cristae morphology that is more similar to that of red-blooded notothenioid species (Johnston et al. 1998). The underlying genome and architecture associated with temperate adaptation in this species is completely unknown. Given that cryonotothenioids share a temperate common ancestor (Matschiner et al. 2011; Near et al. 2012), and other polar-to-temperate transitions have also occurred in several red-blooded cryonotothenioid lineages (Stankovic et al. 2002; Dornburg et al. 2017), understanding adaptive processes active in *C. esox* will uncover whether a single or a diversity of adaptive processes enabled the re-colonization of temperate environments in this group.

To explore the genomic signatures of secondarily temperate adaptation in the pike icefish, we sequenced the genome of both *C. esox* as well as its Antarctic sister species *C. gunnari* using PacBio continuous long read (CLR) sequencing, and generated high quality, chromosome-level genome assemblies for whole-genome comparative analyses. In addition to *C. gunnari*, we included comparisons to six available notothenioid genomes – two icefish, three red-blooded species across two different families, and one temperate outgroup to the Antarctic clade – to detect *C. esox*-specific changes in genome architecture and traits putatively under selection. In addition to the genome assemblies, we generated measures of population-level genetic diversity and divergence for *C. esox* and *C. gunnari*, which we used to uncover patterns of divergence and selection between Antarctic and temperate icefish populations. We found that the *C. esox* genome is largely conserved in structure and organization to that of other characterized icefish genomes; however, we also found evidence of structural variation specific to the *C. esox* lineage. These structural variants often co-occur in genomic regions with elevated genetic divergence and directional selection, highlighting the complex genetic architecture underlying secondarily temperate adaptation. We observe several genes under selection related to the mitochondria and visual perception, suggesting possible mechanisms for adapting to temperate environments subsequent to the specialization to Antarctic cold and oxygen-rich conditions, and unique polar light-dark regime.

## Results

### Genome assembly and annotation

In order to characterize the genetics underlying temperate adaptation in an icefish, we first sequenced the genomes of both the secondarily temperate icefish, *C. esox*, and its Antarctic sister species, *C. gunnari*, using PacBio CLR and scaffolded using Hi-C chromosome conformation capture libraries, generating chromosome-level reference genome assemblies for each species. For *C. esox*, a single male individual from Puerto Natales, Chile (Fig. 1) was used for genome sequencing. The resulting genome assembly was 987 megabasepairs (Mbp) in length (Table 1), comparable with the assembly size of around 1 gigabasepairs (Gbp) described in other icefish genome assemblies (Bargelloni et al. 2019; Kim et al. 2019; Bista et al. 2022; Lu et al. 2022). The scaffold and contig N50 were 43.6 Mbp and 2.6 Mbp, respectively, and 97.72% of the total bases were assembled into 24 putative chromosomes. The final assembly was 96.3% gene-complete, according to analysis by BUSCO v5.1.3, using the Actinopterygii reference gene set. In addition, we sequenced RNA extracted and pooled from several different tissues, which we used as transcript-level evidence for annotating the genome using the BRAKER software (Brůna et al. 2021; Gabriel et al. 2021). A total of 28,257 protein coding genes were annotated in this *C. esox* assembly.

**Table 1:** Genome assembly and annotation statistics.

|  | *C. esox* |  | *C. gunnari* |  |
| --- | --- | --- | --- | --- |
| Genome Assembly |  |  |  |  |
| Scale | Contig | Scaffold | Contig | Scaffold |
| Total Size (bp) | 986,435,181 | 987,108,782 | 993,506,282 | 994,201,109 |
| Number of Fragments | 3,499 | 2,067 | 2,992 | 1,536 |
| Largest Fragment (bp) | 9,841,188 | 56,719,351 | 15,571,738 | 55,323,500 |
| N50 (bp) | 2,611,138 | 43,613,322 | 3,182,242 | 44,089,792 |
| L50 | 107 | 11 | 84 | 11 |
| Number of Chromosome-Scale Scaffolds | -- | 24 | -- | 24 |
| Total Bases in Chromosomes (bp) | -- | 964,606,985 | -- | 978,689,797 |
| Percent of Assembly in Chromosomes | -- | 97.72% | -- | 98.44% |
| Gene Completeness (*BUSCO* v5.1.3) |  |  |  |  |
| Complete | 3,506 (96.3%) |  | 3,506 (96.3%) |  |
| Complete & Single Copy | 3,463 (95.1%) |  | 3,469 (95.3%) |  |
| Complete & Duplicated | 43 (1.2%) |  | 37 (1.0%) |  |
| Fragmented | 14 (0.4%) |  | 19 (0.5%) |  |
| Missing | 120 (3.3%) |  | 115 (3.2%) |  |
| Total | 3,640 |  | 3,640 |  |
| Reference Gene Set | actinopterygii_odb10 |  | actinopterygii_odb10 |  |
| Genome Annotation |  |  |  |  |
| Number of Protein Coding Genes | 28,257 |  | 27,918 |  |
| Number of Exons | 246,908 |  | 249,138 |  |

The *C. gunnari* reference assembly, obtained from a single male individual collected from the West Antarctic Peninsula (Fig. 1) was 994 Mbp in length, with a scaffold and contig N50 of 44.1 Mbp and 3.2 Mbp, respectively (Table 1). A total of 98.4% of bases were assembled into 24 chromosome-level scaffolds, consistent with a haploid chromosome number of 24 by cytogenic characterization of the *C. gunnari* karyotype (Ozouf-Costaz et al. 1996). According to BUSCO V5.1.3, this assembly was 96.3% gene-complete. A total of 27,918 protein-coding genes were annotated based on transcript-level evidence obtained from RNAseq data. For both the *C. esox* and *C. gunnari* assemblies, a complimentary assessment of gene completeness at the protein-level and genome-level is available on (Table S1).

### Conserved synteny between temperate and Antarctic icefish genomes

We performed a conserved synteny analysis using Synolog (Catchen et al. 2009; Small et al. 2016) to identify patterns of genomic organization between the genomes of *C. esox, C. gunnari*, and that of other available, high-quality notothenioid assemblies (Kim et al. 2019; Jo et al. 2021; Bista et al. 2022) (Table S2). In addition to size and chromosome number, the *C. esox* and *C. gunnari* genomes display conservation in large-scale genome organization, showing one-to-one correspondence between 24 orthologous chromosomes (Fig. 2A). Similar patterns can be observed between *C. esox* and other chromosome-scale icefish genomes (Fig. S1-2), demonstrating that the conservation in genome structure in *C. esox* is not specific to the *Champsocephalus* lineage and might be conserved across icefish. However, several examples of intra-chromosomal rearrangements were observed between the two *Champsocephalus* genomes (Fig. 2B-C), with 12 out of the 24 chromosomes displaying genomic rearrangements of considerable length (>1 Mbp). Since we had applied a two-assembly approach for each genome, these chromosomal rearrangements were validated by looking at the patterns of within-species synteny (see Methods). By comparing conserved synteny against the chromosome scale genome assembly of an outgroup, the non-Antarctic notothenioid *Eleginops maclovinus* – the closest sister to the Antarctic clade (Cheng *et al*., *in prep*), we observed examples of rearrangements specific to the *C. esox* lineage, originating in or during its split from the Antarctic *C. gunnari* around 1.6 Mya (Stankovic et al. 2002; Near et al. 2012; Dornburg et al. 2017). One remarkable example is chromosome 21 in *C. esox* (Fig. 2C), which displays a double chromosomal inversion when compared to both the icefish sister species *C. gunnari* and the non-Antarctic *E. maclovinus*.

**Figure 2.**
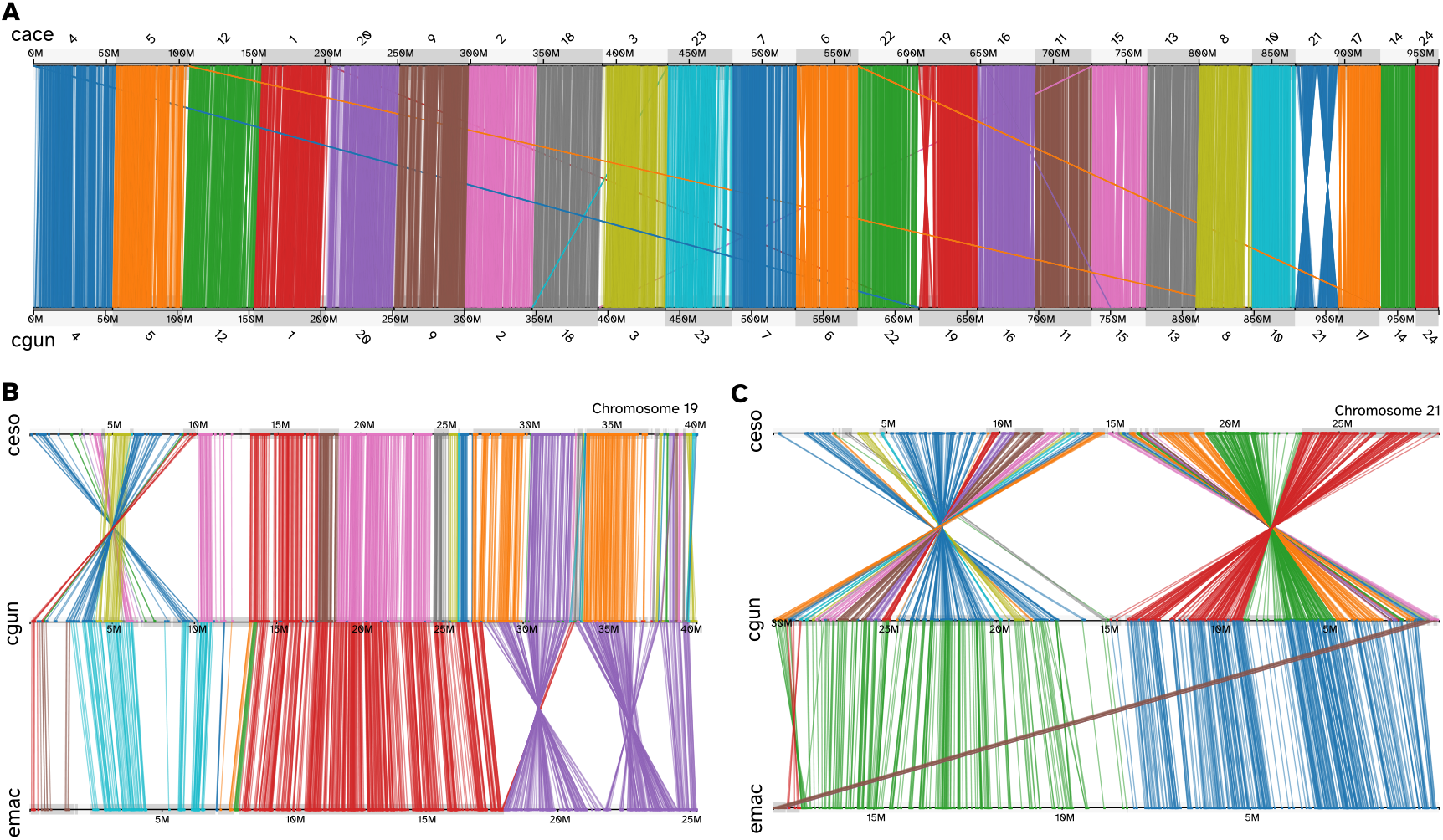
Patterns of conserved synteny between the genomes of temperate and Antarctic icefish species. A) Genome-wide synteny plot shows one-to-one correspondence between the 24 chromosomes of *C. esox* (top) and *C. gunnari* (bottom). Each line represents an orthologous gene between the two genomes, color-coded according to the chromosome of origin. B) Conserved synteny between chromosome 19 in *C. esox* (top), *C. gunnari* (middle), and *E. maclovinus* (bottom). Lines are color-coded according to the genomic contig of origin in either *C. esox* or *E. maclovinus*. A *C. esox*-specific chromosomal inversion is seen in the chromosome, located between 1-10 Mbp of the *C. esox* chromosome. There are two additional inversions between the icefish and *E. maclovinus* lineages. C) Conserved synteny between chromosome 21 in *C. esox* (top), *C. gunnari* (middle), and *E. maclovinus* (bottom). This chromosome displays two large chromosomal inversion specific to the *C. esox* lineage.

### Repeat distributions along the genome

Repeat elements have been associated with patterns of genome evolution in cryonotothenioids, including an expansion in genome size and chromosomal diversification (Detrich et al. 2010; Auvinet et al. 2018; Chen et al. 2019). Therefore, we performed a *de novo* annotation of the repeat elements present in our assemblies to identify changes in repeat content putatively linked to temperate adaptation. Both the *C. esox* and *C. gunnari* genomes appear to contain nearly identical repeat distributions, composed of 59.38% and 59.45% repeat content, respectively (Fig. 3A, Table S3). Highly similar values were also observed for the major interspersed repeat element types, *i.e*., DNA, SINE, LINE, and LTR (Fig. 3A, Table S3). These repeat distributions match those observed in both other icefish genomes (Kim et al. 2019; Bista et al. 2022), and red-blooded Antarctic notothenioids (Jo et al. 2021; Bista et al. 2022) (Fig. 3A), suggesting that repeats in *C. esox* are largely not related to the secondarily temperate adaptation process and might instead be present across species of the Antarctic clade.

**Figure 3.**
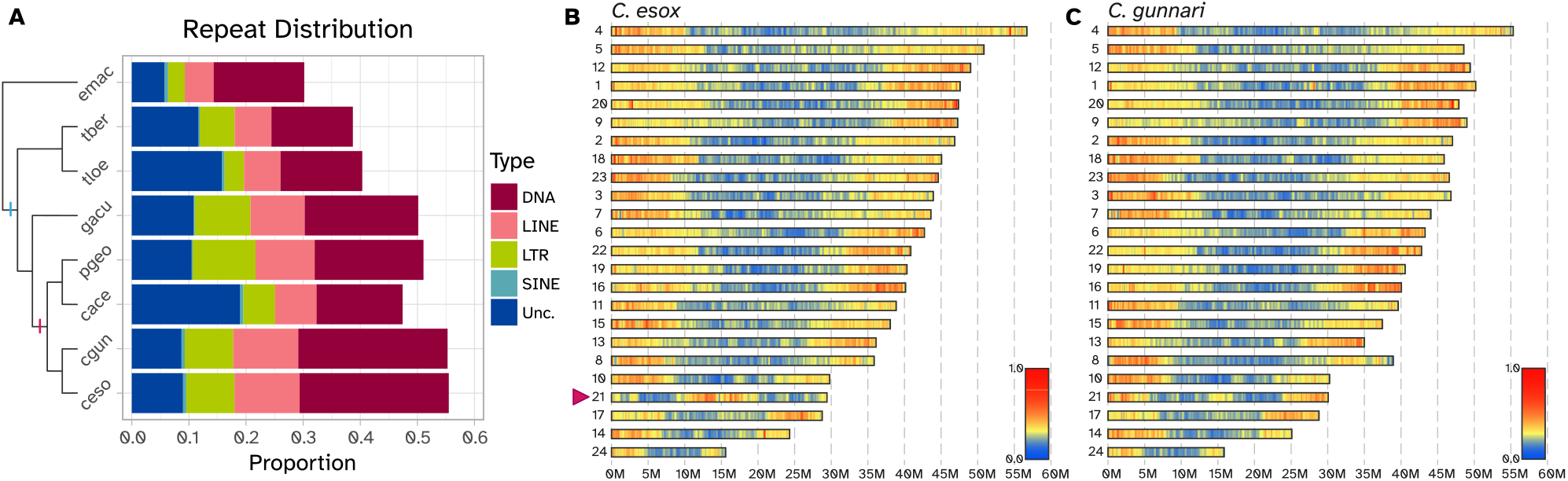
Repeat distribution is conserved along icefish genomes. A) Interspersed repeat distributions for *C. esox* (ceso), *C. gunnari* (cgun), *Chaenocephalus aceratus* (cace), *Pseudochaenichthys georgianus* (pgeo), *Gymnodraco acuticeps* (gacu), *Trematomus loennbergii* (tloe), *Trematomus bernacchii* (tber), and *E. maclovinus* (emac). The cladogram on the left shows the phylogenetic relationship among the species. Blue hatch in the cladogram represents the location of Cryonotothenia, while the red hatch shows the location of Channichthyidae. The colors in the bars represent proportions across five different repeat classes (Unc. = Unclassified). The repeat distributions for *C. gunnari* and *C. esox* were generated in this study. We used the published repeat distributions for the remaining species: *E. maclovinus* (Cheng *et al*., *in prep*), *T. loennbergii* (Jo et al. 2021), *C. aceratus* (Kim et al. 2019), *P. georgianus*, *G. acuticeps*, and *T. bernacchii* (Bista et al. 2022). B) Relative distribution of interspersed repeats along the *C. esox* genome. Chromosomes are ordered according to their size, largest to smallest. The color gradient represents the relative repeat content, plotted for each 250 Kbp window along the genome. Repeats appear to have a higher distribution along the terminal ends of chromosomes, except for chromosome 21 (red arrow) which shows evidence of genomic rearrangements in the lineage. C) Relative distribution of interspersed repeats along the *C. gunnari* genome. Chromosomes are ordered according to the length of the orthologous chromosome in *C. esox*. The color gradient represents the relative repeat plotted for each 250 Kbp window. As in *C. esox*, higher repeat distribution is observed in the terminal ends of chromosomes.

Both *C. esox* and *C. gunnari* display repeats that are more concentrated along the termini of chromosomes than in their centers (Fig. 3B-C). One clear exception is the high repeat content in the center of chromosome 21 in *C. esox* (Fig. 3B), where a species-specific rearrangement is found (Fig. 2C). The double inversion pattern observed would cause the two repeat-rich terminal ends in the *C. gunnari* chromosome to be located in the center of the *C. esox* chromosome.

### Genome-wide diversity and divergence

To measure patterns of genetic diversity and divergence between *C. esox* and *C. gunnari* populations we genotyped individuals using Restriction site-Associated DNA sequencing (RADseq). For each species, we collected individuals across two sites (Fig. 1; hereafter referred to as populations); 28 individuals for *C. esox* (16 from Puerto Natales and 12 from Canal Bárbara), and 68 for *C. gunnari* (48 from South Georgia Island and 20 from the West Antarctic Peninsula). Sequenced reads from these individuals were aligned to our new *C. esox* assembly and genotyped using Stacks v2 (Rochette et al. 2019). After filtering missing data we retained ~56K RAD loci (average locus length of 639 bp) along the genome, comprised of 288K variant sites. The two *C. gunnari* populations, from South Georgia Island and the West Antarctic Peninsula, were more divergent from one another than the two populations of *C. esox* from Puerto Natales and Canal Bárbara (Table S4). To limit the effects of this within-species population structure, we removed the West Antarctic Peninsula *C. gunnari* samples from later steps of the analysis, assaying *C. esox* samples in a single, species-level population (see Methods). At this species-level comparison, we observe higher genetic diversity in the *C. gunnari* populations (variant-site *π* of 0.12) than in the *C. esox* populations (*π* of 0.088). We also find high genetic divergence between *C. esox* and *C. gunnari* populations (mean *F_ST_*=0.40, mean *D_XY_*=0.0042). Given the high proportion of elevated *F_ST_* values observed between the populations of both species (Fig. S3), likely due to differentially fixed sites resulting from a recent history of isolation, we decided to use the absolute divergence as a comparative metric for subsequent analysis. Regions of high genetic divergence were spread across the genome, with 23 out of 24 chromosomes containing at least one genomic window with outlier *D_XY_* values.

Patterns of elevated genetic divergence between populations can be the product of both selective and neutral processes. To further test the patterns of selection behind regions of genetic divergence, we calculated extended haplotype homozygosity (*EHH*, (Sabeti et al. 2002)) separately in each species. The *EHH* statistic looks for signatures of positive selection in a population by identifying changes in homozygosity and linkage disequilibrium across the genome. We then compared the patterns of *EHH* between *C. esox* and *C. gunnari* populations using the cross-population *EHH* statistic (*XP-EHH*, (Sabeti et al. 2007)), which allowed us to identify changes in *EHH* – and thus differences in positive selection – between temperate and Antarctic icefish populations. Patterns of population-level directional selection appeared to be more common in *C. esox* than in *C. gunnari*, spanning all 24 chromosomes. Out of the 976 sites displaying outlier *XP-EHH* values, 939 (96.2%) showed directional positive selection in temperate populations. There was also strong co-localization between patterns of divergence and selection, as windows of divergence (*D_XY_*) and *EHH* outliers overlapped in 20 chromosomes. Furthermore, there was a correlation with the presence of genomic rearrangements and these selection/divergence outliers (Fig. 4). While not all selection/divergence outlier windows were colocalized with structural differences – and not all structural differences display divergence and/or selection outliers – there were examples of these outlier windows limited to the boundaries of a genomic inversion, *e.g*., chromosome 12 (Fig. 4A), or outlier windows located within the span of a larger chromosomal rearrangement, *e.g*., chromosome 21 (Fig. 4B).

**Figure 4:**
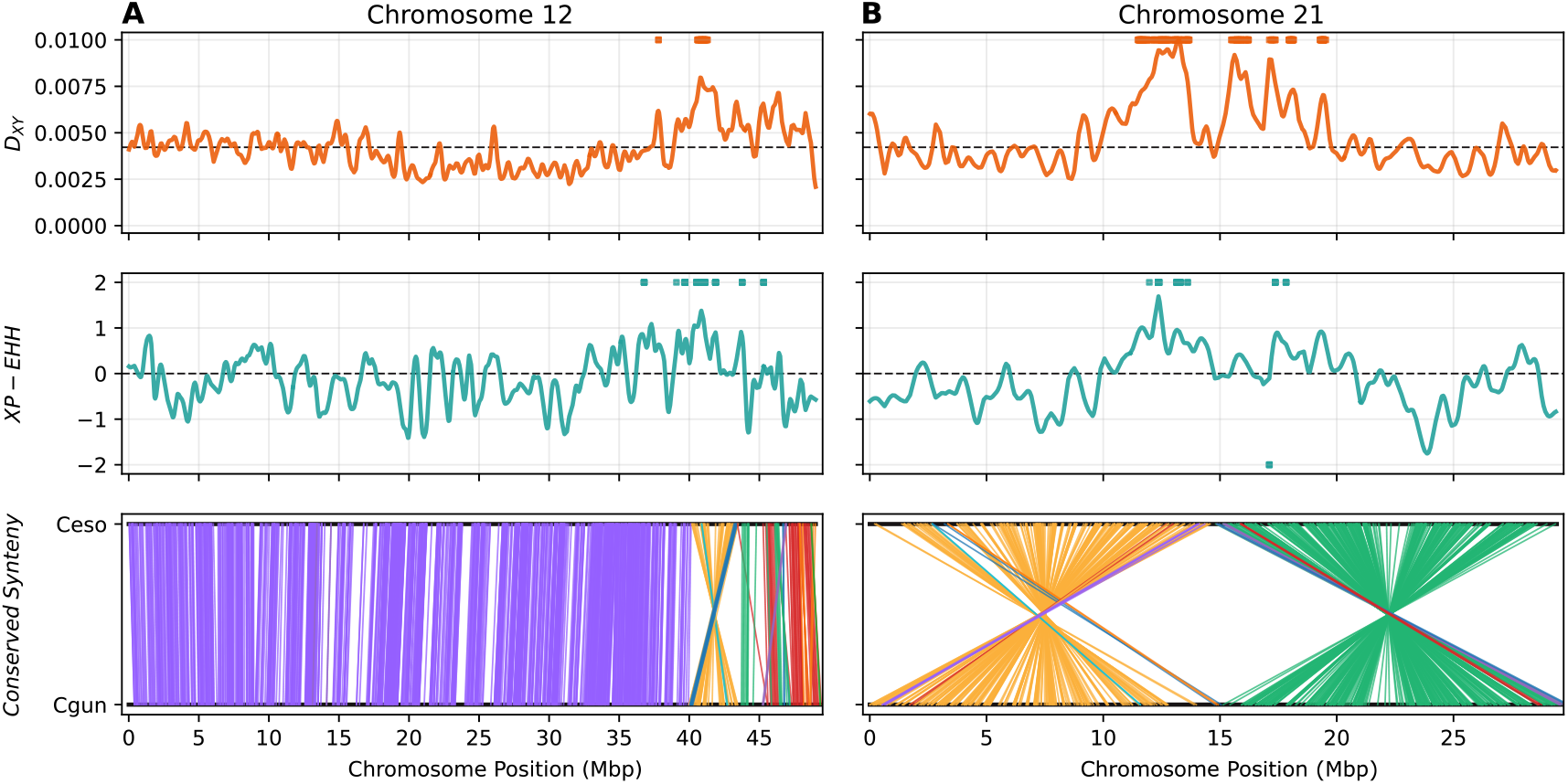
Patterns of genetic divergence and positive selection in populations of temperate and Antarctic icefish species. A) Population-level comparisons for *C. esox* chromosome 12. Top panel shows the distribution of absolute genetic divergence (*D_XY_*) between *C. esox* and *C. gunnari* populations. Orange line shows the kernel-smoothed *D_XY_* values. Top horizontal bars indicate genomic windows displaying outlier genomic divergence (*p* < 5×10^-4^). Middle panel shows the distribution of *XP-EHH* values between *C. esox* and *C. gunnari* populations. Positive values indicate positive selection in *C. esox*, negative values selection in *C. gunnari*. Cyan line shows the kernel-smoothed *XP-EHH*. Horizontal bars indicate variant sites displaying outlier *XP-EHH* values (FDR < 5%). Bottom panel shows patterns of conserved synteny between the *C. esox* and *C. gunnari* orthologous chromosome. Each line represents an orthologous gene, color coded according to its conserved synteny cluster. In chromosome 12, we can observe a region (at 40-45 Mbp) displaying outlier divergence and *XP-EHH* values coinciding with a chromosomal inversion. B) Population level comparisons for *C. esox* chromosome 21. Top, middle, and bottom panels show *D_XY_*, *XP-EHH*, and conserved synteny, as described in panel A. This chromosome displays two large chromosomal inversions (one from 0 to15 Mbp, second one from 15 to 30 Mbp), each co-localized with regions with outlier *D_XY_* and *XP-EHH* values.

### Candidate genes under selection

We used a two-pronged approach to identify candidate genes under selection in *C. esox*. First, we applied an analysis of molecular evolution using the branch-site model implemented in the software PAML (Yang 2007), to identify significant differences in the rate of non-synonymous to synonymous substitutions (*ω*) in the protein-coding genes of *C. esox* relative to orthologs of five other cryonotothenioids. With this approach, we identified a set of genes displaying elevated *ω* values, a measure of positive selection in the focal lineage (Fig. S4). By comparing the annotations of six notothenioid taxa, including four icefish and two red-blooded Antarctic species, we identified 14,093 orthogroups composed of one single-copy-ortholog per-species. Out of these 14K orthogroups, 258 genes displayed significantly elevated *ω* values (hereafter referred to as *dN/dS* candidates, Table S5).

We also identified protein-coding genes that are located within regions displaying population-level signatures of selection, as measured by *XP-EHH* and genetic divergence. This approach found 1,490 protein-coding genes, hereafter referred to as ‘divergence/*XP-EHH* candidates’ (Table S6). While not all these genes have substitutions in their coding sequences, we corroborated the presence of nearby SNPs with outlier *EHH* values, which may reflect cis-regulatory selection (Fig. S5).

The two analyses yielded 1,744 genes under selection representing candidates for secondarily temperate adaptation in *C. esox*. Assessing the functional annotation of these candidates, obtained during genome annotation by comparing the gene sequences to the InterProScan database (Quevillon et al. 2005), we observed genes associated with mitochondrial morphology and maintenance, genes associated with the mitochondrial electron transport chain, with light sensing and vision, and with the heat shock response (Table 2). A complete list of all identified candidate genes and their predicted function based on their zebrafish orthologs is presented in Table S7.

**Table 2:**
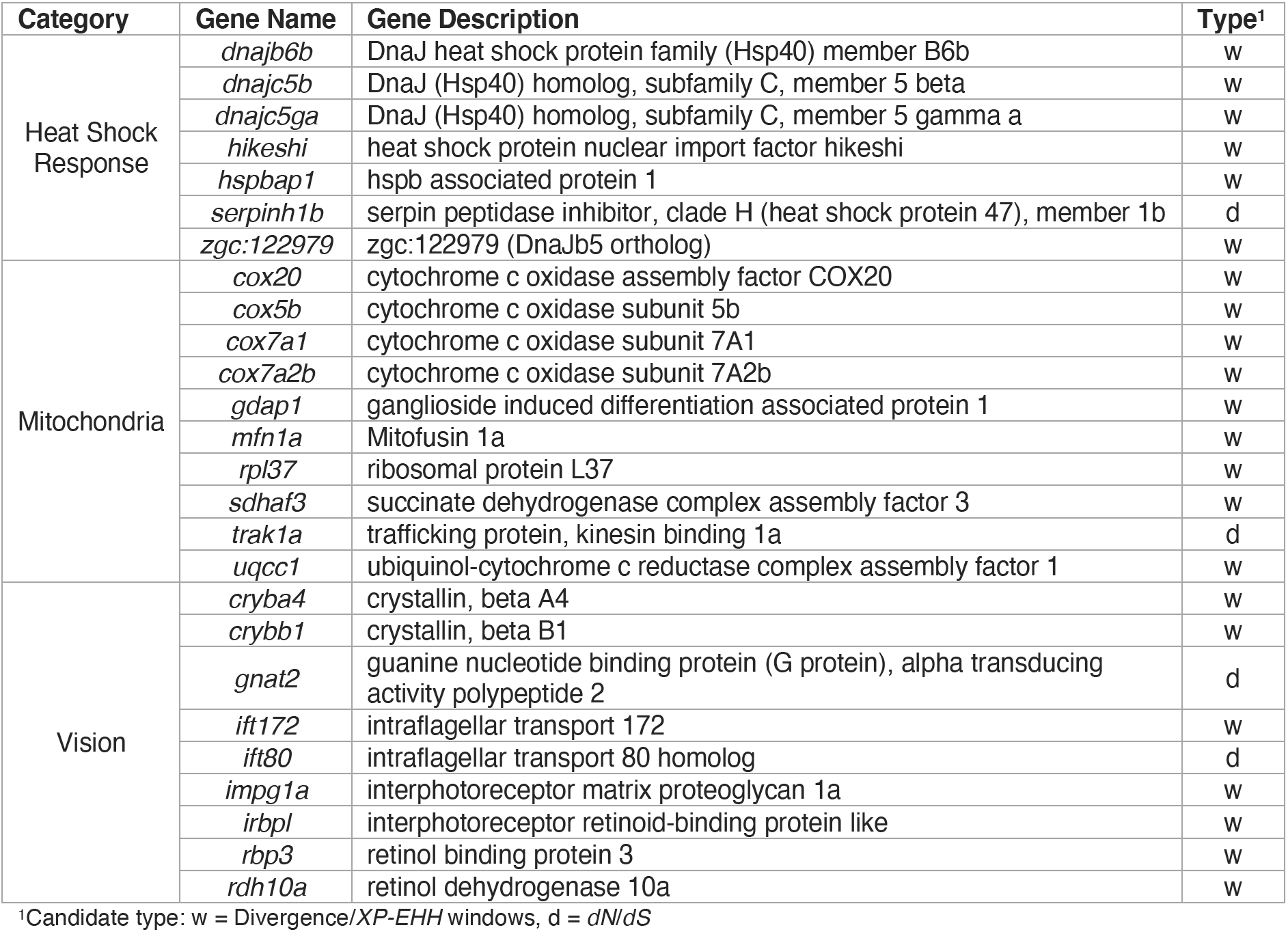
Examples of biological categories displaying top candidates under selection.

### Expanded and contracted gene families

In addition to specific genes exhibiting signatures of positive or directional selection in *C. esox* we identified gene families showing elevated rates of evolution based on the gain and loss of genes within the family. Across seven notothenioids species – six cryonotothenioids including *C. esox* and a temperate outgroup (see Methods), the OrthoFinder software (Emms and Kelly 2015; Emms and Kelly 2019) identified 26,838 phylogenetic hierarchical orthogroups or gene families. CAFE v5 (Mendes et al. 2021) retained 21,019 families that were present at the root of the phylogeny of the sampled species and could be used to infer gene family expansion and contraction events. Out of these 21K gene families, 6,011 exhibit significant events of expansions and/or contractions across one or several points of the species phylogeny (Fig S6). In the temperate pike icefish genome in particular, we observe 495 gene family expansion and 1,080 contraction events (1,575 total). Among the significant expanded/contracted gene families in *C. esox* (Table S8), we observe the contraction of *zona pellucida*, odorant receptor, and PR/SET domain-containing genes, as well as the expansion of several myosin H families, histone methyltransferases, and BED zinc finger protein genes.

### Evolution of antifreeze glycoproteins

The evolution of AFGPs has been regarded as the key evolutionary innovation that allowed notothenioids to survive in the inhospitable Antarctic environment. The cryonotothenioid *AFGP* genotype evolved once (Chen et al. 1997), and its genomic location and gene neighborhood appear to be conserved in a “canonical” *AFGP/TLP* locus across characterized species (Nicodemus-Johnson et al. 2011; Kim et al. 2019; Bista et al. 2022). Given the secondary escape of the *C. esox* lineage into a temperate environment, studying its *AFGP* genotype contributes to understanding gene evolution following the loss of a selective constraint. Even after the temperate transition, *C. esox* displays conservation in the organization of the canonical *AFGP/TLP* locus flanked by *hsl* and *tomm40*, on its 5’- and 3’-end respectively, as in other cryonotothenioids (Fig. 5). The locus contains 5 tandem duplicates of *tryp1*, one copy each of *tryp3* and *TLP*, and six *AFGP* copies. Five of the *AFGP* copies appear nonfunctional (denoted with *Ψ*, Fig. 5). Two of these are partial copies composed of only exon 2, containing the coding sequence of the repetitive AFGP tripeptides (Thr-Ala/Pro-Ala)_n_. The missing exon 1 encodes the signal peptide, thus these partial copies lack an appropriate translation start and the signal sequence for extracellular export of the protein. The other three *AFGP* copies exhibit inactivating frameshift mutations and/or premature stop codons. This high number of pseudogenized *AFGP* copies (5 of 6) likely reflects a history of relaxed selective pressure following secondary colonization of temperate habitats in this lineage. The canonical *AFGP/TLP* locus in the Antarctic sister species *C. gunnari* exhibits the same conserved organization, with variation in the number of copies and direction of several genes. There are eight tandemly duplicated *tryp1* copies, typical of other notothenioids including the basal *E. maclovinus*, versus five in *C. esox*. Three functional *AFGP* copies are observed in the canonical *AFGP/TLP* locus of *C. gunnari* along with one additional, non-functional copy.

**Figure 5:**
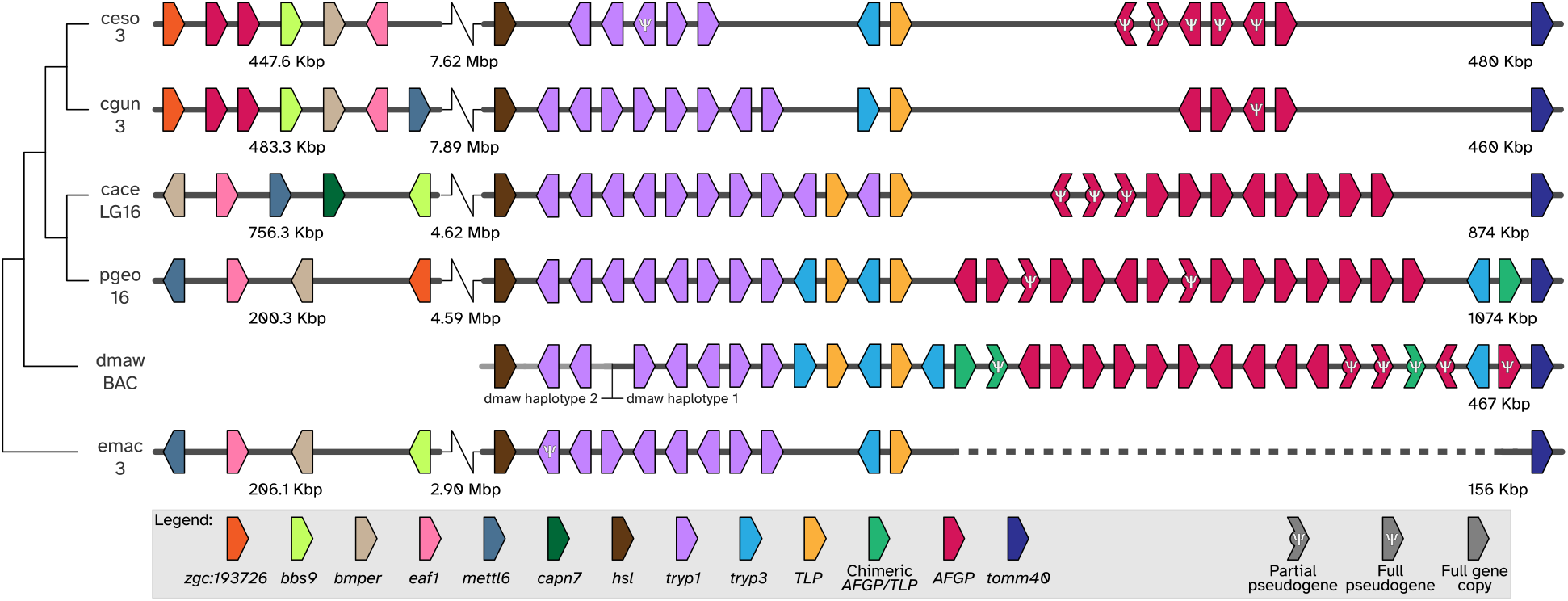
Evolution of the antifreeze glycoprotein locus. Conserved synteny in the *AFGP/TLP* locus for six notothenioids (ceso, *C. esox*; cgun, *C. gunnari*; cace, *C. aceratus*; pgeo, *P. georgianus*; dmaw, *D. mawsoni*; emac, *E. maclovinus*). Cladogram on the left shows the phylogenetic relationship across the six species. The ID for the chromosome containing the *AFGP* is displayed under the species ID. Each horizontal grey line represents the span of the chromosome, arranged from 5’ to 3’ orientation, with the diagonal line representing a break in the represented sequence. For *D. mawsoni*, the sequence represents the bacterial artificial chromosome (BAC) sequence assembled in (Nicodemus-Johnson et al. 2011), with the majority of the sequence (from the third *tryp1* copy to *tomm40*) corresponding to haplotype 1. The horizontal light grey line on the 5’-end of this sequence, from *hsl* to the second *tryp1* copy, corresponds to the 5’-end boundary of the *D. mawsoni AFGP/TLP* haplotype 2. Each colored arrow represents a different gene, color-coded according to their corresponding gene family, and arranged according to their relative position in the annotated sequence. Distances between genes are not drawn to scale. Pentagonal arrows represent full gene copies, while truncated arrows represent partial copies. The Greek letter *Ψ* denotes pseudogenes. Both *C. esox* and *C. gunnari* have relative conservation in the organization of the canonical *AFGP*/*TLP* locus (flanked by the genes *hsl* and *tomm40*) when compared to other Antarctic notothenioid fishes, including both white-blooded (*C. aceratus*, *P. georgianus*) and red-blooded (*D. mawsoni*) species, but display a reduction in *AFGP* gene copies in the canonical locus. *C. esox* additionally shows an increase in pseudogenized *AFGP* copies, with 5 out of the 6 genes appearing nonfunctional. Both *Champsocephalus* species display a small non-canonical locus with two complete *AFGP* copies in the same chromosome but about 8 Mbp away from the canonical locus. Gene synteny in this non-canonical locus is highly conserved between *C. gunnari* and *C. esox*, with both *AFGP* copies flanked by the genes *zgc:193726* and *bbs9*. While the majority of flanking genes (*e.g*., *bbs9*, *bmper*, *eaf1*) are observed across all six species, changes in gene order and orientation are observed between both *Champsocephalus* species and other notothenioids, suggesting ancestral rearrangements in this section of the chromosome.

Of the six notothenioids we examined, excluding the non-Antarctic outgroup *E. maclovinus* (which lacks *AFGPs), C. gunnari* and *C. esox* had the fewest *AFGP* copies. However, in the genomes of both species we found two complete and (at a sequence-level) putatively functional *AFGP* copies about 8 Mbp away from the canonical locus on the same chromosome. In both *C. esox* and *C. gunnari*, these additional copies are flanked by *zgc:193726* (193726 zebrafish transcript homolog) and *bbs9* (Bardet-Biedl syndrome 9), on the 5’- and 3’-ends respectively. These two additional *AFGP* copies have not been identified in any notothenioid species studied to date and, in combination with their conserved synteny across both genomes, might reflect a translocation forming a non-canonical *AFGP* locus unique to the genus *Champsocephalus*.

## Discussion

In this study, we describe two high quality, chromosome-scale genome assemblies for a pair of sister icefish species, the temperate *C. esox* and the Antarctic *C. gunnari*. This species pair represents the only example of major ecological divergence in the Channichthyidae – two closely related species occurring in disparate environments across the Antarctic Polar Front. Their close evolutionary relationship presents a unique opportunity for studying the genetic basis of secondary adaptation to warm environments, which would otherwise be lethal to the cold-specialized cryonotothenioids. The hemoglobin-less phenotype precluding active transport of oxygen in icefishes makes them particularly vulnerable to warm temperatures, in turn making *C. esox* an even more remarkable example of adaptation and diversification from a cold-specialized and stenothermal ancestor.

### Conservation in genome structure despite ecological disparity

We observed a high degree of conservation in genome structure and organization in the temperate *C. esox* and Antarctic *C. gunnari* genomes, despite drastic ecological disparity and specialization. Both genomes show conservation in architecture and chromosome number, including a strong one-to-one correspondence across orthologous chromosomes. These results are in agreement with the 2n=48 karyotype previously characterized in *C. gunnari* (Ozouf-Costaz et al. 1996) and the patterns of conservation in chromosome number and organization described in other cryonotothenioid genome assemblies (Kim et al. 2019; Bista et al. 2020; Bista et al. 2022). While changes in chromosome number have been observed in cryonotothenioids (Ozouf-Costaz et al. 1991; Morescalchi et al. 1992; Ghigliotti et al. 2015) – some of them quite extreme, as seen in the genus *Notothenia* (Amores et al. 2017) – the ancestral teleost karyotype number (N=24) is largely conserved. However, other changes to genome organization, like changes in karyotype structure, have been extensively observed (Mazzei et al. 2006; Ghigliotti et al. 2007; Ghigliotti et al. 2015), giving precedence to the patterns of intra-chromosomal changes observed between *C. esox* and *C. gunnari*.

The distribution of transposable repeat elements is also conserved between temperate and Antarctic icefish genomes. This distribution has long been thought to be important to genome evolution and lineage-specific diversity in vertebrates (Warren et al. 2015). Genomic repeats comprise a large portion of cryonotothenioid genomes (Auvinet et al. 2018; Kim et al. 2019; Bista et al. 2022), with a lineage-specific expansion in transposable elements likely linked to environmental stressors in the extreme Antarctic environment. Note that, while the percentage of the genome composed of transposable elements is higher for the two *Champsocephalus* species when compared to other published notothenioid assemblies (Fig. 3), this is likely a technical artifact explained by methodological differences. For example, when we recomputed the repeat annotation for *Chaenocephalus aceratus* (Kim et al. 2019) using the analyses described in this study (see Methods), the repeat distribution increased to that observed in the two *Champsocephalus* assemblies (Fig. S7). This technical issue aside, given the known link between environmental stress and the expansion of transposable elements (Oliver and Greene 2009; Lanciano and Mirouze 2018), it is striking that no substantive change in the transposable element distribution is observed in *C. esox*. The transition to a temperate environment following extreme cold specialization, both in terms of novel environmental stressors and possible demographic changes, could have otherwise been predicted as a prime driver for transposable element expansion and diversification. However, the relatively recent divergence time between *C. esox* and the Antarctic icefish lineage (~1.6 Mya (Stankovic et al. 2002; Near et al. 2012; Dornburg et al. 2017)) may not be sufficient for extensive diversification, resulting in a genome that still retains many of its ancestral Antarctic properties and organization. This highlights the need for the study of other secondarily temperate notothenioid genomes, particularly for species exhibiting more ancient polar-to-temperate transitions, to determine if the patterns described above are specific to *C. esox* or shared across independent polar-to-temperate transition events.

### The genomic architecture of secondarily temperate adaptation

Colonization of temperate Patagonian waters by the ancestral *C. esox* lineage likely occurred over the last 2 million years (Stankovic et al. 2002), matching the Quaternary colonization of temperate environments observed in other red-blooded notothenioid species (Hüne et al. 2015; Ceballos et al. 2019). Given this relatively recent window of time, it is remarkable to find a complex architecture underlying polar-to-temperate transitions in this system, as it would require the organization and interaction of many individual loci for adaptation to take place. We identified divergence outlier loci present in most *C. esox* chromosomes (Fig. S3), indicating high sequence differentiation between temperate and Antarctic *Champsocephalus* populations. This fact alone is unsurprising since these two species are separated by the Antarctic Polar Front, which serves as a strong barrier for gene flow between population residing in the waters of Patagonia and the Southern Ocean (Poulin et al. 2014; Moore et al. 2018). However, neutral processes cannot fully account for the divergence as many of these regions also house candidate genes under selection, signaling that changes in selective pressures during the polar-to-temperate transition acted across the whole genome and, in part, mediated the observed patterns of divergence.

Genomic rearrangements are likely playing an important role in shaping the genomic architecture underlying this adaptive process. We identified several examples of genomic rearrangements between the *C. esox* and *C. gunnari* genomes co-localized with the candidate regions described above (Fig. 4). Structural variation, particularly chromosomal inversions, are mediators of genomic processes underlying adaptation, preventing recombination between inverted chromosomes, and maintaining divergent adaptive haplotypes (reviewed in (Kirkpatrick and Barton 2006; Wellenreuther et al. 2019)). Chromosomal inversions have played a role in several adaptive processes in wild populations, including thermal adaptation in *Drosophila* (Puig Giribets et al. 2019), local climatic adaptation in seaweed flies (Mérot et al. 2021), the maintenance of social morphs in *Formica* ants (Brelsford et al. 2020), differentiation between migratory and non-migratory Atlantic cod populations (Matschiner et al. 2022), and divergence between coat color and tail length in deer mice ecotypes (Hager et al. 2022). Observing these candidates of selection within inversions implies that these structural rearrangements are acting (or acted in the past) as islands of divergence, allowing for the maintenance and segregation between cold- and temperate-adapted haplotypes in the emerging *C. esox* populations after their isolation from cold-specialized populations. This would suggest that the structural variants, and the regions contained within that gave rise to temperate-adapted haplotypes, may have existed in the Antarctic icefish population prior to speciation. These two factors of the genomic architecture – structural variants co-located with signatures of selection alongside high baseline differentiation in the absence of gene flow – illustrates a contrast between early and late evolutionary forces. Instead of the classic pattern of divergence with gene flow, *i.e*., a handful of regions in the genome displaying elevated patterns of divergence surrounded by a low divergence background (Cruickshank and Hahn 2014), based on our data, we hypothesize that early adaptive divergence was facilitated by structural variants, after which neutral variation independently accumulated between Antarctic and temperate populations following isolation due to geographic and environmental barriers. Following this early isolation and the processes of temperate adaptation, we may be presently observing the remnants of these ancient islands of divergence on this now differentially adapted and fully speciated icefish populations.

The key role of inversions as islands of divergence is primarily observed in other systems as polymorphic variants in a single population or multiple populations in sympatry or with some degree of gene flow. While our reconstructions of single genomes cannot show if these inversions were (or still are) polymorphic in either *C. gunnari* or *C. esox*, the overlap in the genome between structural variation and candidates of selection suggests a possible link between the two. Whether they served as mechanisms of isolation in the emerging populations or facilitated the maintenance of novel adaptive haplotypes following speciation, these chromosomal inversions are a key component of the genomic architecture of secondarily temperate adaptation and speciation in *C. esox*. In order to fully understand their role, however, future studies should focus on determining both the polymorphic status of these rearrangements, as well as determining their age in relation to the time of speciation between *C. esox* and *C. gunnari*.

### Physiological and molecular changes in a polar-to-temperate transition

The process of adaptation to stable temperature environments has been experimentally shown to limit physiological plasticity in response to future changes to those temperatures (Morgan et al. 2022). Thus, the ability of a cold-specialized icefish species to adapt to temperate environments is a remarkable example of adaptation following specialization. Given the candidate genes we identified in *C. esox*, we can observe how secondarily temperate adaptation has impacted an already unique icefish physiology.

#### Antifreeze glycoproteins

Previous work detected the presence of AFGP coding sequences in *C. esox* (Miya et al. 2016), but it was unknown if they were functional. Our genome assemblies demonstrate that, in the absence of chronic cold, the *AFGP* copies in *C. esox* have followed a different evolutionary trajectory from those in the Antarctic notothenioids. A canonical *AFGP/TLP* locus remains in *C. esox* (Fig. 5), which is not unexpected given the recent age of divergence from its Antarctic ancestor. Yet, while similar in copy number, the majority of *C. esox AFGP* coding sequences exhibit evidence of pseudogenization, showing a combination of exon loss, premature stop codons, or frameshift mutations that would prevent the production of AFGP polyproteins. While pseudogenization of *AFGP* copies has occurred in both red- and white-blooded Antarctic species (Fig. 5), the particularly high frequency in *C. esox* likely resulted from the absence of freezing-related selective pressures in their temperate environment. Interestingly, the sequence of three *AFGP* copies appear to remain complete and intact (one in the canonical locus, and two in the novel, translocated locus). However, our preliminary results indicate absence of functional AFGPs in *C. esox* (unpublished), and the mechanisms of AFGP trait loss in this system remain under investigation.

In notothenioids, the AFGP gene is known to have evolved once from a trypsinogen-like protease (*TLP*) precursor (Chen et al. 1997), and the extant gene family has a conserved gene neighborhood across different cryonotothenioid families (Nicodemus-Johnson et al. 2011; Kim et al. 2019; Bista et al. 2022). Variation in copy number appears to evolve via tandem duplications within the *AFGP/TLP* locus. Strikingly, our results show that both *C. esox* and *C. gunnari* have two additional *AFGP* copies translocated outside the canonical site (Fig. 5), a feature not observed in all characterized notothenioids to date. This non-canonical *AFGP* locus displays conserved synteny between the two species (and the locus placement is more broadly supported by conserved synteny of the surrounding gene neighborhood of more basal taxa), suggesting that the translocation predates speciation and originated in the Antarctic ancestor of both *C. esox* and *C. gunnari*. Interestingly, we find evidence of additional, small-scale rearrangements in the chromosome containing the canonical and novel *AFGP* loci, both when comparing *C. gunnari* to *C. esox* and to other species (Fig. 5, Fig. S8). While none of these changes disrupt the *AFGP/TLP* locus, their presence suggests chromosomal volatility. Previous studies have described the presence of polymorphic *AFGP* loci in cryonotothenioids, with different haplotypes displaying copy number variation in *AFGP* and surrounding genes (Nicodemus-Johnson et al. 2011). Therefore, future studies on this non-canonical *AFGP* locus should explore the presence of polymorphic *AFGP* haplotypes in *Champsocephalus*, particularly with respect to these additional copies. For example, determining if the non-canonical locus is polymorphic itself or if it is linked to a specific *AFGP*/*TLP* haplotype.

#### Mitochondria

White-blooded icefishes display divergent mitochondrial form and function when compared to other red-blooded notothenioid species (O’Brien and Mueller 2010), arising from evolutionary changes of the organelle to compensate at least for the loss of hemoglobin (Sidell and O’Brien 2006; Bargelloni et al. 2019). Previous studies described *C. esox* as possessing higher mitochondrial density than other Antarctic icefish (Johnston et al. 1998) and having cristae (*i.e*., inner membrane) morphology more similar to that of red-blooded notothenioids (Johnston et al. 1998; O’Brien and Mueller 2010). In addition, recent analysis of the *C. esox* mitochondrial genome identified unique patterns of gene duplication and the likely presence of heteroplasmy in the species (Minhas et al. 2022). Correlating with these previously described changes, we found several candidate genes under selection that are related to mitochondrial morphology, potentially mediating the unique mitochondria observed in this species. The gene *mfn1a* encodes for a GTPase essential for mitochondrial fusion (Chen et al. 2003), while *gdap1* is known to affect several mitochondrial phenotypes including changes to fragmentation and elongation (Niemann et al. 2005). The interaction between the gene products of *mfn1a* and *gdap1* are known to maintain mitochondrial morphology and stability across the cellular mitochondrial network (Niemann et al. 2005). In addition, the gene *trak1a*, for which we found non-synonymous substitutions under positive selection in *C. esox* (Table S5), plays a role in mitochondrial trafficking (Koutsopoulos et al. 2010). The trak1 protein has been found to colocalize with mitofusin proteins, like those encoded by *mfn1a*, putatively facilitating mitochondrial fusion (Lee et al. 2018).

Aside from mitochondrial morphology, we identified other candidate genes under selection in *C. esox* related to cellular respiration and the organization of the electron transport chain. More specifically, we identified several genes that form or are associated with cytochrome *c* oxidase (COX), *i.e*., complex IV of the electron transport chain. Two genes that appeared within our divergence/*XP-EHH* windows (Table S6) are *cox7a1* and *cox7a2b*, two paralogs of COX subunit 7. Each paralog produces a different, tissue-specific protein isoform (reviewed in (Sinkler et al. 2017)), which, in mammals, appears to fine tune complex IV activity via the formation of different COX dimer complexes (Zong et al. 2018). Another gene within this set of candidates is *cox5b*, which encodes for a hypoxia-inducible isoform of COX subunit 5 (Hodge et al. 1989; Kwast et al. 1998). While not a component of complex IV itself, *cox20* is another candidate of selection involved in the assembly and maturation of the COX supermolecule (Bourens et al. 2014). In addition to complex IV, we also identified other related genes as putative candidates of selection. For example, the ribosomal protein *rpl37* (located in one of the large inversions on chromosome 21) plays a role in the assembly of complexes I and III of the electron transport chain (reviewed in (Soto et al. 2012)). The gene *sdhaf3* is important for the maturation of complex II (Na et al. 2014), while *uqcc1* acts on the assembly of complex III (Brockerhoff et al. 2003). While none of these candidate genes contain non-synonymous substitutions that could be altering their canonical function, mutations over nearby non-coding elements could alter their regulation and expression patterns. The supramolecular organization of the oxidative phosphorylation molecular complexes is dynamic. Different molecular assemblages of the different electron transport chain complexes can be found across different tissue types, oxygen levels, and resource conditions (reviewed in (Enríquez 2016)). Therefore, temperate adaptation in *C. esox* may have acted upon the regulation of components of the electron transport chain, taking advantage of the variability in the configuration of this complexes to maintain or maximize metabolic output in its new warmer, less oxygenated environment.

#### Circadian rhythms and vision

One prior study reported the loss of genes belonging to the canonical circadian regulation in cryonotothenioids (particularly, in the blackfin icefish *C. aceratus*), likely as a result of the unique polar light-dark regimes experienced by Antarctic organisms (Kim et al. 2019). We find that many of the same canonical circadian regulation genes are also apparently lost in *C. esox* (Fig. S9), including the absence of several paralogs of the period (*e.g*., *per1a*, *per2a*, *per3*) and cryptochrome circadian regulators (*e.g*., *cry1b*, *cry2*, *cry3b*, *cry4*). Many of these putative loses are in fact shared between *C. esox*, its Antarctic sister species *C. gunnari*, and other cryonotothenioids (Fig. S9, Table S9), implying that they are shared across the lineage and not a product of the temperate adaptation process. The molecular and physiological changes resulting from these gene losses in icefish are presently unknown, as well as their interaction with other abiotic factors (*e.g*., light, temperature) that might be interacting with biological rhythms in these organisms. Particularly, given the complex patterns of loss across circadian regulator paralogs across teleosts (Toloza-Villalobos et al. 2015), these loses alone are likely not sufficient to indicate a disruption in the canonical circadian regulation in these organisms. We observe a single circadian-related candidate gene under selection in *C. esox*, a serotonin receptor (*htr7a*). While this gene is not part of the canonical circadian regulation pathways, the neurotransmitter serotonin does exhibit circadian rhythmicity (Morin 1999), and might thus indicate some yet uncharacterized changes to circadian-associated pathways in *C. esox*.

Changes in light patterns can impact the biological clocks of organisms (Beale et al. 2013). *C. esox* originated from a cold-specialized ancestor that might have been subjected to polar regimes of extended months of light and dark periods in the Antarctic. It is important to note that *C. gunnari* has an extensive range and can be found on both the Atlantic and Indian sectors of the Southern Ocean (Kock and Everson 2003). Within the Atlantic sector, *C. gunnari* can be found along a 10-15 degrees longitudinal range, from the West Antarctic Peninsula in the south, to the Scotia Sea in the north (Fig. 1). South Georgia island, located at the northernmost point of this distribution, is at a similar latitude to that of Patagonia, and thus would have similar periods of daylight (Smith and Walton 1975). This creates environmental differences among these different *C. gunnari* populations, which likely include differences in daily light levels. In turn, these different *C. gunnari* populations might experience different selective pressures related to photoperiod and circadian rhythms. However, while the patterns of population structure and isolation among these *C. gunnari* populations has been studied (Kock and Everson 2003; Damerau et al. 2014; Young et al. 2015), their evolutionary relationship to extant *C. esox* populations remain unknown.

While the light environment *C. esox* experiences may or may not resemble that of other closely related Antarctic icefishes, we do find evidence of selection acting on genes related to light sensing and vision. Among our candidates in divergence/*XP-EHH* windows (Table S6) we found two crystallin genes, *cryba4* and *crybb1*, both encoding for β-crystallin proteins. Crystallins are integral structural components of the vertebrate eye lenses (reviewed in (Wistow et al. 2005; Wistow 2012)), whose development is mediated by the complex regulation of crystallin genes across cell types and developmental stages (Farnsworth et al. 2021). Compared to other teleosts, Antarctic notothenioids possess cold-stable crystallin proteins that remain transparent at subzero temperatures (Kiss et al. 2004). It is unknown how the lens of temperate notothenioids compares to the Antarctic species. However, changes in the regulation of *cryba4* and *crybb1* in *C. esox* might allow for the development of functional lenses at higher temperatures, counteracting the effects of the cold specialized phenotypes observed in other notothenioids.

In addition to structural components of the lens, we observed several candidates under selection related to visual processes. One example among the *dN/dS* candidates (Table S5) is *gnat2*, which encodes the alpha subunit (GNAT2) of a cone-specific transducin (Morris and Fong 1993), a heterotrimeric G protein coupled to the photoreceptor in the visual phototransduction cascade (reviewed in (Arshavsky et al. 2002)). Transducin is activated by a light-activated opsin, whereby its GNAT2 subunit binds a GTP, dissociates from the heterotrimer and activates the next component – a phosphodiesterase. The codon in *gnat2* under selection in *C. esox* is a Thr, which substitutes a Gly that is conserved across most of the background notothenioid group (Fig. S10). This substitution suggests a novel protein phenotype present in the secondarily temperate *C. esox*; however, the functional outcome of this substitution in the signaling activity of *C. esox* GNAT2 awaits further exploration. Two genes among our *divergence/XP-EHH* window candidates are *irbpl* and *rbp3*, both interphotoreceptor binding protein homologs that are important for the transport of visual pigments in the light cycle (Jin et al. 2009; Parker et al. 2011). Another candidate, *impg1a*, encodes a proteoglycan protein part of the interphotoreceptor matrix, an extracellular structure that plays a role in the maintenance of photoreceptor cells and cellular adhesion to the retinal pigmented epithelium (Ishikawa et al. 2015). In *Astyanax* cavefish, *impg1a* is among the differentially expressed genes associated with loss of vision (Stahl and Gross 2017), although the functional role of *impg1a* on this phenotype is unknown. Additionally, we observed two intraflagellar transport genes, *ift172* and *ift80*, as candidates of selection. Intraflagellar transport proteins are essential for flagellar assembly, which then has an ubiquitous role in the transport of a variety of biomolecules (reviewed in (Rosenbaum and Witman 2002)). Both *ift172* and *ift80* have been specifically described as important for the transport of opsin molecules in zebrafish (Sukumaran and Perkins 2009; Hudak et al. 2010). Lastly, another gene located in the chromosome 21 inversion and among our list of divergence/*XP-EHH* window candidates is *rdh10a*. In mice, the enzyme encoded by this gene plays a role in the regeneration of 11-*cis*-retinal, an important reaction in the recycling of retinoids during the visual cycle (Sahu et al. 2015).

Our candidate genes do not intrinsically indicate differences in the circadian rhythm and biological periodicity between *C. esox* and other cryonotothenioid species. However, we find evidence for selection in the maintenance, transport, and activation of visual pigments and tissue. While the genetic differences described above could simply be related to sensing in its light environment, detection of light alone can be important for the entrainment of the internal biological clock of the organism to its environment (Hunter-Ensor et al. 1996; Hurd and Cahill 2002). The presence and detection of light plays a key role upstream of the core circadian rhythm molecular feedback loop in zebrafish (Vatine et al. 2011). Therefore, changes in light detection, as a result of selection in these vision-related genes, could lead to them playing a larger role in the entrainment of biological rhythms in the temperate pike icefish.

#### Heat shock response and molecular chaperones

Associated with cold specialization in cryonotothenioids is their inability to mount the classic, inducible heat shock response (HSR) (Hofmann et al. 2000; Bilyk and Cheng 2014; Bilyk et al. 2018). Instead, increased constitutive expression of *hsp6a*, the major ancestrally inducible member of the HSP70 family, is observed across both red- and white-blooded Antarctic species, suggesting an increase in native expression to mitigate cold denaturation of proteins (Bilyk et al. 2021). In a studied icefish (*Chionodraco rastrospinosus*), exposure to thermal stress triggers an inflammatory response instead of the inducible HSR (Bilyk et al. 2018). *C. esox* lives in an environment that can reach temperatures of ~14°C in the austral summer, so biological pathways related to stress-induced chaperones and HSR represent prime candidates for selection. Among our divergence/*XP-EHH* window candidates (Table S6) we observed a suite of cytosolic molecular chaperones, among them several DnaJ/HSP40 paralogs, including *dnajc5b*, *dnaj5ga*, *dnajb6b*, and *zgc:122979* (a *dnajb5* homolog). DnaJ/HSP40 are a family of molecular co-chaperones that bind to and stabilize HSP70s and other chaperones (reviewed in (Qiu et al. 2006)). Furthermore, we also identified *hspbap1*, which encodes a HSP27-associated protein. HSP27s are a suite of small HSP molecular chaperones that have been associated with responses to oxidative stress (reviewed in (Arrigo 2001)). Another HSR-associated gene found in divergence/*XP-EHH* windows is *hikeshi*, a HSP70 nuclear import carrier important for HSR gene transcription and the reversal of stress induced phenotypes (Kose et al. 2012).

Lastly, we found evidence of non-synonymous substitutions under selection in *serpinh1b*, which produces an inducible procollagen-specific HSP47. Unlike other molecular chaperones, HSP47 recognizes the folded state of its target, the collagen triple helix (Widmer et al. 2012). Expression of *serpinh1b* has been associated with thermal stress response in the rainbow trout (Wang et al. 2016; Pandey et al. 2021). While most of these genes do not show coding changes under selection, mutations over nearby non-coding sites might be driving the evolution of HSR regulation. The state of inducible HSR in *C. esox* is presently unknown; however, evidence of selection on these genes suggests possible changes in HSR pathways and other chaperones following colonization of temperate environments and are good candidates for future work regarding the functional characterization of secondarily temperate adaptation.

## Conclusion

Here, we describe new genomic resources for *C. esox*, the only non-Antarctic icefish, and its Antarctic congener, *C. gunnari*. We characterize the primary changes to the genomic architecture of this temperate icefish, including several major structural variants, and show where key signals of evolutionary forces, including measures of divergence, linkage, and genic selection co-localize with these physical, chromosomal changes. In addition, we identified genes displaying evidence of positive selection in the species, providing examples of major biological and cellular pathways likely underlying the physiological changes behind secondary adaptation to temperate environments. These lines of evidence paint a picture of how *C. esox* has evolved since its Antarctic departure, allowing for a polar-to-temperate transition in this group and illuminating an important evolutionary question of how adaptation can occur after specialization. Future studies should explore the genotypes and candidate genes identified here in more detail, measuring tissue-specific gene expression of these genes, characterizing the effect of any nonsynonymous mutations altering protein function, and complimenting these results with the physiological studies of live individuals. Nonetheless, the work presented here sets the stage for future projects to further research the mechanisms behind secondarily temperate adaptation in the cryonotothenioid system.

## Materials and Methods

### Specimens and sampling

A total of 28 *C. esox* and 68 *C. gunnari* individuals were collected and used in this study (Fig. 1). Twelve of the 28 *C. esox* were captured with a gill net from Canal Bárbara (Fig. 1 – CB), a channel connecting the Strait of Magellan to the Pacific Ocean, in April 2016. The other 16 were obtained by rod and reel or gill net in the Patagonia waters near Puerto Natales, Chile (Fig. 1 – PN) in 2008, 2009, 2016, and from December 2017 to early January 2018. *C. gunnari* was collected from two Antarctic regions. Forty-eight of the 68 were collected from waters around South Georgia Island and Shag Rock Shelf (Fig. 1 – SG) by trawls from onboard the British Antarctic Survey chartered Falkland Islands F/V *Sil* (ZDLR1) during the South Georgia groundfish survey from January – February 2017. The other 20 individuals were collected from several coastal sites along the West Antarctic Peninsula (Fig. 1 – AP) using otter trawl from onboard the US R/V Laurence M. Gould during austral winters (July – August) of 2008 and 2014. Detailed metadata for the specimens collected and used for the various sequencing effort in this study are given in Table S10.

For genome sequencing, white muscles from a single male *C. gunnari* (collected in 2014 from the Gerlache Strait in West Antarctic Peninsula) were flash frozen with liquid nitrogen and stored at −80°C until used for high molecular weight (HMW) DNA extraction. For *C. esox*, fresh liver from a single male (collected in January 2018 from Puerto Natales) was perfused to obtain isolated hepatocytes. Perfusion was carried out on ice with collagenase (Sigma) at 0.5 mg/mL in heparinized, Ca^+2^-free notothenioid physiological buffer (260 mM NaCl, 5 mM KCl, 2.5 mM MgCl_2_, 2.5 mM NaHCO_3_, 2 mM NaH_2_PO_4_, 250 units heparin mL^-1^, pH 8.4), following a published protocol (O’Grady et al. 1982). Cell density was assessed by counting with a hemacytometer, and appropriate volumes of the hepatocyte suspension were embedded in 1% agarose blocks using disposable plug molds (BioRad). Hepatocytes were thoroughly lysed in-agarose using 1% lithium dodecyl sulfate in notothenioid ringer (100 mM sodium phosphate buffer, pH 8.0, adjusted to 420 mOsm with NaCl). The blocks were then fully equilibrated with a preservation buffer (100 mM EDTA, 10 mM Tris, 0.2% N-laurylsarcosyl, pH 9.0) that permitted return transport at ambient temperature, and were subsequently kept at 4°C until use. For RAD and RNA sequencing, tissue samples for both species were dissected, preserved in ice-cold or −20°C 90% ethanol, and stored at −20°C until use.

Specimen collection and sampling were conducted with permits granted by the Chilean Fisheries Service under Technical Memorandum P.INV N° 244-2016, the Government of South Georgia & the South Sandwich Islands Regulated Activity Permit 2016-026, and the University of Illinois at Urbana-Champaign IACUC approved protocols 07053 and 17148, for the respective field collection and sampling effort.

### HMW DNA and long-read genome sequencing

HMW DNA of the single male *C. gunnari* was extracted from frozen white muscle of the single male using the Nanobind Tissue Big DNA Kit (Circulomics) following manufacturer protocol. For *C. esox*, agarose blocks of embedded hepatocytes from the single male were treated with proteinase K (final concentration 800 *μ*g/*μ*L, 52°C, 2 hr.). The treated blocks were then melted (65°C, 20 min.), and digested with 2 units of β-agarase (NEB) per block (1 hr., 42°C). The liberated HMW DNA was gently extracted once with phenol-chloroform (1:1) and precipitated with isopropanol. The recovered DNA was further purified using the Nanobind Tissue Big DNA Kit following manufacturer protocol starting at the DNA binding step. Quality check of HMW DNA included fluorometric (Qubit v.3, Invitrogen) quantification of concentration, spectrophotometric (Nanodrop One, Fisher Scientific) determination of purity, and HMW assessment using pulsed field electrophoresis (CHEF Mapper XA, BioRad), and Fragment Analyzer (Advanced Analytical).

Pacific Biosciences CLR library preparation and sequencing were carried out at the University of Oregon Genomics & Cell Characterization Core Facility. The HMW DNA was lightly sheared with Megaruptor (Diagenode) at 60 kilobasepairs (Kbp) target length for library construction using PacBio SMRTbell Express Template Prep Kit 2.0. The resulting library was selected for inserts approximately greater than 30 Kbp with the BluePippin (Sage Science) and sequenced on two SMRT cells 8M for each species on Sequel II for 30 hours of data capture. A total of 12.6 million and 10.7 million CLR reads were obtained for *C. esox* and *C. gunnari* respectively. Statistics of sequence reads are given in Table S11.

### Hi-C library and sequencing

For scaffolding the genome assembly, Hi-C chromosome conformation capture libraries were constructed for the same individual of *C. esox* and *C. gunnari* using ethanol-preserved spleen and liver respectively. Library constructions were performed by Phase Genomics, Inc. using its proprietary Proximo Hi-C kit that utilizes *DpnII* as the restriction enzyme. The libraries were sequenced by Phase Genomics on an Illumina NovaSeq6000, which yielded 257.8 million and 161.2 million 2×150 bp reads for *C. esox* and *C. gunnari*, respectively.

### Transcriptome library and sequencing

RNAseq libraries for *C. esox* and *C. gunnari* were constructed and sequenced to provide transcript-level evidence for annotating protein-coding genes in the genome. RNA from a broad range of tissues (14 for *C. esox*, 19 for *C. gunnari*) derived from multiple individuals were isolated using Ultraspec II (Biotecx) or Trizol (Invitrogen) reagent, DNaseI treated, purified, and checked for RNA quality on Agilent Bioanalyzer. RNA from all tissues achieved a high RNA Quality Number (RQN), as shown in Table S12. The individual, tissue-specific RNA samples from each species were then pooled at 1*μ*g or 0.5 *μ*g to form a final pool. The library was then constructed from 2 *μ*g of the final pool using the TruSeq Stranded mRNAseq Sample Prep Kit (Illumina). The two libraries were multiplexed and sequenced for 2×250 bp reads on one Illumina HiSeq2500 lane. Raw reads for *C. esox* were processed for quality using Trimmomatic v0.32 (Bolger et al. 2014), retaining 98.7 million reads-pairs. For *C. gunnari*, raw reads were independently processed using AfterQC v0.9.6 (Chen et al. 2017), leading to 99.9 million read-pairs retained.

### RADseq library and sequencing

To genotype *C. esox* and *C. gunnari* individuals, a Restriction site-Associated DNA (RADseq) library was constructed. Genomic DNA from the 28 *C. esox* and 68 *C. gunnari* individuals were isolated from ethanol-preserved muscle or other tissues using standard tissue lysis and phenol-chloroform extraction protocol. DNA concentrations were measured using a Qubit fluorometer (Invitrogen). A single enzyme digest RADseq library was constructed following published protocols (Baird et al. 2008; Etter et al. 2011) with slight modifications. Briefly, the DNA of the 96 individuals was diluted to equimolar concentration (33.3 g/*μ*L), and 1*μ*g (30 *μ*L) of each sample was transferred to and digested in a 96-well plate with the restriction enzyme *SbfI*-HF (NEB). Custom P1 adapters (Hohenlohe et al. 2012), each containing a 7 bp unique barcode identifier, were then ligated to the digested DNA of each individual. The 96 barcoded samples were pooled at equimolar concentration for a total of 2 *μ*g per pool to generate four replicate pools. Each pool was sheared using a Covaris M220 focused ultrasonicator, after which the fragmented DNA was size-selected within a 300-600 bp range using AMPure XP beads (Beckman Coulter). The size-selected pools were then end-repaired and A-tailed, followed by ligation of P2 adapters, forming the library. To minimize PCR duplicates, replicate libraries were pooled to maximize the amount of template DNA amplified, pooling ~150 ng of template DNA per replicate. A 100 *μ*L PCR reaction was used to amplify the samples for 12 PCR cycles. The final amplified library was cleaned with AMPure XP beads and quantified using Qubit fluorometry. The library was sequenced on an Illumina NovaSeq6000 SP 2×150 bp lane at the University of Illinois Roy J. Carver Biotechnology Center, producing 821 million raw reads.

### Genome Assembly

PacBio CLR raw reads were subsampled at different depths of coverage and size distributions (*i.e*., removing reads shorter or longer than a specified range of length). This subsampling of reads served as an assembly optimization process (Rayamajhi et al. 2022). See supplements for more details on the subsampling optimization. For *C. esox*, the highest quality assembly was derived from a subsample of reads 10-50 Kbp in length that provided 70× coverage, based on an initial genome size estimation of 1.1 gigabasepairs (Gbp), yielding 3.8 million reads with a 20.4 Kbp mean length and 23 Kbp read N50. For *C. gunnari*, the most optimal subsample contained reads over 15 Kbp and 80× coverage, yielding a 29.5 Kbp mean length and a 32 Kbp N50. Contig-level assemblies were made using two different assemblers, Flye (Kolmogorov et al. 2019) and wtdbg2 (Ruan and Li 2020). Each assembly was assessed for contiguity using QUAST v4.4 (Gurevich et al. 2013) and gene completeness using BUSCO v3.0.1 (Simão et al. 2015), using the `actinopterygii_odb9` lineage dataset. For both species, Flye v2.5 generated superior contig-level assemblies based on quality and completeness scores (Supplements – Methods) and was selected as the primary assembly for downstream analyses. The wtdbg2 v2.5 assembly is referred to as the secondary assembly and was used for comparison purposes (Table S13).

The Hi-C data were then integrated into both the primary and secondary assemblies to generate chromosome-level super-scaffolds using Juicer v1.6.2 (Durand et al. 2016) to identify the Hi-C junctions, and then with Juicer’s `3d-dna` program to complete the integration of scaffolds. In addition, conserved synteny analysis (described below) between primary and secondary assemblies was used to perform manual curation of the scaffolding process. For example, we employed manual insertions, inversions, or translocations of whole contigs when discrepancies were found between the primary and secondary assemblies, and evidence, such as contig or scaffold boundaries, that supported one assembly to be correct. We used a custom Python program to propagate these changes through the constituent assembly files, such as the structure (AGP), annotation (GFF), and sequence files (FASTA). Following scaffolding and manual curation, a re-assessment of the contiguity of the final assembly was done using Quast v4.4 and an assessment of gene-completeness was performed with both BUSCO v3.0.1 and BUSCO v5.1.3 (Manni et al. 2021), using the `actinopterygii_odb9` and `actinopterygii_odb10` gene datasets, respectively.

### Genome Annotation

To annotate repeat elements in the genome, a *de novo* repeat library was built for each reference assembly separately using the RepeatModeler v2.0.2a pipeline (Flynn et al. 2020). First, a database was generated for each genome with `BuildDatabase`, using the NCBI database as input (`-engine ncbi`). The final library was then created using `RepeatModeler`, enabling the optional discovery of LTR elements using LTRharvest (`-LTRStruct`) (Ellinghaus et al. 2008). The *de novo*, species-specific repeat libraries were then combined with the teleost-specific repeat libraries available in Repbase release 27.02 (Bao et al. 2015). This combined repeat library was then used as the input of RepeatMasker v4.1.2-p1 (Smit et al. 2013), which was used for the final repeat annotation and masking of the genome assemblies.

For the annotation of protein-coding genes, we first indexed the masked reference genome for alignment using STAR v2.7.1.a (Dobin et al. 2013) `--runMode genomeGenerate`, after which we aligned the processed RNAseq reads to the chromosome-level genome assembly using `--runMode alignReads`. Final annotation of protein-coding genes was done using TSEBRA (Gabriel et al. 2021), which selects curated transcripts generated by the BRAKER annotation pipeline (Hoff et al. 2016; Brůna et al. 2021). For integration with TSEBRA, we ran BRAKER v2.1.6 in both protein and transcript modes. For protein mode (`--prot_seq`), we used the zebrafish protein sequences from OrthoDB v10.1 (Kriventseva et al. 2019), while the transcript mode (`--bam`) used the species-specific RNAseq alignments. The output of both BRAKER runs was then processed using TSEBRA v1.0.1 to retain the annotation of protein coding-genes supported at both the protein and transcript level. For both species, we verified the gene-completeness of the annotation using BUSCO v5.1.3, providing the amino acid sequences of the annotated protein-coding genes as input, running in protein mode (`--mod prot`) and comparing against the `actinopterygii_odb10` reference set. Finally, the functional annotation of the curated protein-coding genes was done using the InterProScan database (Quevillon et al. 2005).

For both *C. esox* and *C. gunnari*, BRAKER failed to annotate the AFGP genes. Thus, genomic coordinates and sequences for all *AFGP* copies were determined by manual annotation. We used `blastn` from BLAST+ v2.4.0 (Camacho et al. 2009) to locate the coding sequences of both *AFGP* and other genes in the canonical *AFGP*/*TLP* locus (Nicodemus-Johnson et al. 2011), including trypsinogen 1 (*tryp1*), trypsinogen 3 (*tryp3*), trypsinogen-like protease (*TLP*), chimeric *AFGP/TLP* genes, and the two flanking genes, *hsl* (hormone sensitive lipase, *lipeb* homolog) and *tomm40* (translocase of outer mitochondrial membrane 40). For *AFGP* and *AFGP/TLP* sequences, we used the *Dissostichus mawsoni*-specific orthologs (Nicodemus-Johnson et al. 2011) as queries in the search. Coordinates of the `blastn` alignments were used to guide manual inspection and mapping of complete gene structures, including intron-exon boundaries, and the precise start and end of the highly repetitive tripeptide (Thr-Ala/Pro-Ala)_n_ *AFGP* coding sequences (Chen et al. 1997). The *AFGP* copies were then translated and manually inspected to evaluate the integrity of the gene, or whether mutations had occurred resulting in pseudogenization. We used comparative synteny (see Methods below) to assess conservation in the position of flanking genes surrounding the *AFGP/TLP* locus. We compared the annotation between both *C. esox* and *C. gunnari*, as well as *C. aceratus* (Kim et al. 2019) and *Pseudochaenichthys georgianus* (Bista et al. 2022), to confirm their location in orthologous chromosomes. In addition, we discovered a non-canonical *AFGP* locus during manual annotation (see Results), which we validated by applying a similar conserved synteny assessment of the surrounding genomic neighborhood between *C. esox* and *C. gunnari*.

Given both the repetitive nature and tandem duplication of *AFGP* copies in notothenioid genomes (Nicodemus-Johnson et al. 2011; Kim et al. 2019; Bista et al. 2022), which could lead to their misassembly, we further validated the regional assembly of the *AFGP/TLP* locus by tiling the raw CLR reads in this portion of the genome. Raw reads were aligned using minimap2 v2.24-r1122 (Li 2018). In addition, we identified *AFGP*-specific k-mers in both the reference sequences and raw reads (Supplements – Methods). We visualized both the read alignments and presence of *AFGP* k-mer “clumps” (series of sequential k-mers) using a custom Python program, confirming that: 1) the read alignments spanned the totality of the corresponding *AFGP* locus, 2) no major disruptions in read alignments were present, and 3) that the location of *AFGP* clumps in the raw reads matched the location of *AFGP* sequences manually annotated in the reference sequences (Fig. S11-14). Further validation of the non-canonical *AFGP* copies was done by inspecting for significant secondary/supplementary alignments between both *AFGP* loci, confirming that raw reads are uniquely mapping to their corresponding *AFGP* locus of origin.

### Conserved synteny analysis

An analysis of conserved synteny was performed at different stages of the assembly using the Synolog software (Catchen et al. 2009; Small et al. 2016). Annotated coding sequences from *C. esox and C. gunnari* primary and secondary assemblies, and other species of interest, were first reciprocally matched using the `blastp` algorithm from BLAST+ v2.4.0. The BLAST results and genome annotation coordinates were fed to Synolog, which 1) establishes reciprocal best BLAST hits to identify orthologous genes, 2) uses the genome coordinates of each best hit to define clusters of conserved synteny between the genomes, 3) refines ortholog assignments using the defined clusters, and 4) defines orthology between chromosomes/scaffolds based on the conserved synteny patterns. In addition to the *C. esox* and *C. gunnari* genomes described here, for comparative purposes, we included other high-quality notothenioid assemblies including *C. aceratus* (Kim et al. 2019), *P. georgianus* (Bista et al. 2022), *Gymnodraco acuticeps* (Bista et al. 2022), and *Trematomus bernacchii* (Bista et al. 2022) (Table S2).

This conserved synteny analysis was initially used to manually curate the assemblies, by identifying discrepancies between primary and secondary assemblies, and/or to identify structural variants located within scaffold boundaries indicating a misassembly. Once the curated chromosome-level sequences were generated, conserved synteny was determined between the genomes of *C. esox* and *C. gunnari* against the genome of *Eleginops maclovinus*, the closest sister species to the Antarctic clade, in order to identify and name orthologous chromosomes. The *E. maclovinus* assembly (Cheng *et al*., *in prep*) was first compared against the *Xiphophorous maculatus* reference assembly, which contains the ancestral teleost karyotype number of 24 chromosomes (Amores et al. 2014). Chromosomes were named accordingly to these patterns of conserved synteny.

### Molecular evolution on protein-coding genes

To identify patterns of molecular evolution and positive selection on protein coding genes, we first identified single-copy orthologous genes (orthogroups) with Synolog, as described above. A total of 14,093 orthogroups were identified across the six notothenioid species (*i.e*., *C. esox*, *C. gunnari*, *C. aceratus*, *P. georgianus*, *G. acuticeps*, and *T. bernacchii*). To guide alignments, we used the published notothenioid phylogenetic tree (Near et al. 2018), using the ape v5.5 R package (Paradis and Schliep 2019) to sample the tree to include only the taxa of interest (Fig. S4). The coding sequences for all genes in each orthogroup were aligned using PRANK v.170427 (Löytynoja and Goldman 2005), using the codon alignment option (`-codon`), a guide phylogenetic tree (`-t`), and forcing insertions to be skipped (`-F`). Alignments were then filtered using Gblocks v 0.91b (Castresana 2000) under a codon model (`-t=c`). We used GWideCodeML v 1.1 (Macías et al. 2020), a wrapper for PAML v4 (Yang 2007), designed for large-scale genomic datasets, to identify genes under positive selection in *C. esox* based on the ratio (*ω*) of non-synonymous (*dN*) to synonymous substitutions (*dS*). GWideCodeML was run under the branch-site model (`-model BS`), supplying a species tree (`-tree`), defining *C. esox* as the foreground species (using the `-branch` file), and generating a *dN/dS* summary for all tested orthogroups (`-dnds`). We used a custom Python script to further parse the GWideCodeML outputs and obtain additional information needed to filter the results (*e.g*., site class proportions, model likelihoods, and per-site posterior probabilities). We then selected orthologs showing *ω* values greater than one, with alignment lengths greater than 150 amino acid sites, and showing significant (*p*-value < 0.05) differences in the likelihood ratio test between alternative and null models. For these selected genes, we also looked for individual sites showing posterior probabilities greater than 95% for *2a* or *2b* site classes. In addition, we filtered comparisons for which both the *2a* and *2b* site class proportions were zero, as these can result in unreliable *ω* estimates (Yang and dos Reis 2011). The gene ontology information for all the retained genes under selection was then categorized using the PANTHER database (Mi et al. 2021).

### Expansion and contraction of gene families

We used the CAFE v5 (Mendes et al. 2021) software to identify evolutionary patterns of gene family expansion and contraction in *C. esox* relative to the other studied cryonotothenioid species. To do this, we first selected the annotated protein sequences for six cryonotothenioids (*C. esox*, *C. gunnari*, *C. aceratus*, *P. georgianus*, *G. acuticeps*, and *T. bernacchii*) and one temperate outgroup (*E. maclovinus*), which were then clustered into orthogroups using the OrthoFinder v2.5.4 software (Emms and Kelly 2015; Emms and Kelly 2019). When running the OrthoFinder pipeline, we first inferred trees based on multiple sequence alignments (`-M msa`). Alignments were made using MAFFT v7.310 (Katoh and Standley 2013), and the final gene trees inferred using FastTree v2.1.11 (Price et al. 2010). Once the orthogroup inference was complete, we used a custom Python script to calculate a per-species tally of the number of genes in each OrthoFinder phylogenetic hierarchical orthogroup. These tallied orthogroups, alongside the pruned phylogenetic tree from (Near et al. 2018), were then used as input for CAFE v5.0.0. As described in the CAFE documentation, we ran the software in three separate iterations in order to 1) estimate the global error model attributed to issues in genome assembly and annotation (`--error_model`) (Han et al. 2013); 2) calculate the mean rate of gene gain and loss (*λ*) accounting for the estimated error (`-e<error_file>`); 3) to perform an estimation of cladewise gene family expansion and contraction using this fixed, error-adjusted *λ* value (`--fixed_lamda`). Orthogroups with significant expansion and/or contraction events were determined according to the *p*-value reported by CAFE (*p* < 0.05).

### RADseq data processing

The 821.4 million paired-end reads from the sequenced library were processed with Stacks (Rochette et al. 2019). Raw reads were analyzed with `process_radtags` v2.54 to demultiplex samples, removing those with low quality (`--quality`), containing uncalled bases (`--clean`), lacking the *SbfI* cutsite (`--renz-1 sbfI`), containing adapters (`--adapter_1, --adapter_2`), and rescuing barcodes and cutsites (`--rescue`). After filtering, 737,838,374 reads (89.8%) were retained, with a mean 7.7 million reads retained per sample. The processed reads were then aligned to the *C. esox* reference assembly generated in this study with BWA `mem` v0.7.17-r1188 (Li 2013), using SAMtools v1.12 (Li et al. 2009) to process and sort alignments.

Following alignment, samples were genotyped using `gstacks` v2.59, removing PCR duplicate reads (`--rm-pcr-duplicates`). After removing 46.5% PCR duplicates, the library contained a non-redundant average coverage of 23.8×. A total of 9 samples were removed due to low coverage (<5×). The resulting 87 samples (24 for *C. esox* and 63 for *C. gunnari*) were processed using the `populations` module, defining the *C. esox* and *C. gunnari* samples as two distinct populations. To filter for missing data, we retained only loci present in the two populations (`-p 2`), in 80% of samples per population (`-r 80`), and variant sites with a minimum allele count of 3 (`--min-mac 3`). The final Stacks catalog kept 56,275 loci and 288,631 variant sites.

### Genetic divergence and signatures of selection

To determine patterns of genetic divergence across the genome between *C. esox* and *C. gunnari*, the Stacks catalog described above was reanalyzed using `populations` v2.60. To account for the effects of genetic divergence between the two *C. gunnari* populations (see Results), the West Antarctic Peninsula individuals were removed from the analysis. The final comparisons for divergence and selection were made between 69 individuals, 24 *C. esox* (combined across Canal Bárbara and Puerto Natales) and 45 *C. gunnari* individuals from South Georgia Island. In addition to filtering for missing data, we calculated *F*-statistics between the populations (`--fstats`) and averaged the resulting statistics along the chromosomes using a kernel-smoothing algorithm (`--smooth`). The `populations` program was run twice using the parameters described above. First, using the `--bootstrap-archive` flag to generate an archive of randomly selected loci. Once the archive was generated, we applied bootstrap resampling 10,000 times over all kernel-smoothed statistics (`--bootstrap, --bootstrap-reps 10000`). Genomic windows displaying outlier values of genetic divergence between *C. esox* and *C. gunnari* populations were identified based on the *p*-values from bootstrap replicates (*p* < 5×10^-4^).

In addition to genetic divergence, this RADseq dataset was used to calculate extended haplotype homozygosity (*EHH*, (Sabeti et al. 2002)) to identify population-level patterns of positive selection. The Stacks catalog was again filtered using `populations`, this time increasing the stringency of filtering parameters to minimize the imputation of missing genotypes during phasing (`-r 90`, `-p 2`, `--min-mac 3`), and exporting the output as a sorted VCF (`--ordered-export`, `--vcf`). This stringent filtering still retained 53,464 loci and 239,008 variant sites. Before phasing, we split the Stacks VCF output into two species-specific VCFs using VCFtools v0.1.15 (Danecek et al. 2011) (`--keep`) in order to phase sites for each species separately. Each species-specific VCF was phased using beagle v5.3 (Browning and Browning 2007; Browning et al. 2018). We provided an initial effective population size of 5,000 (`ne=5000`), based on the *N_e_* estimation for the related blackfin icefish (Kim et al. 2019). Population size and error parameters were estimated (`em=true`) by running for 12 iterations (`niterations=12`). The phased VCF files were then indexed and merged using BCFtools v1.12 (Danecek et al. 2021). *EHH* calculations were performed using the rehh v3.2.1 R package (Gautier et al. 2017). First, the phased VCF was loaded using the data2haplohh() function, creating a subset of the data for each species, and filtering sites with a minimum allele frequency less than 5%. The integrated, site-specific *EHH* (*iES*, (Tang et al. 2007)) was calculated in each species using the scan_hh() function. To account for gaps of ungenotyped sequences in the genome in the RADseq dataset, gaps were scaled (`scalegap`) to 20.3 Kbp (median distance between RAD loci in the dataset), with a max gap (`maxgap`) distance of 104.5 Kbp (95^th^ percentile of the inter-locus distance) and keeping the per-site *EHH* integration in the boundaries of gaps (`discard_integration_at_border=FALSE`). Species-specific *iES* values were compared using the cross-population *EHH* statistic (*XP-EHH*, (Sabeti et al. 2007)) using the `ies2xpehh()` function, obtaining both *XP-EHH* and a significance value (-log10(*p*-value)) at each site. To account the effects of multiple testing, a Benjamini-Hochberg false discovery rate (FDR) correction was applied to the *XP-EHH* significance *p*-value using the `p.adjust()` function in R v4.0.1 (R Core Team 2020). Sites having corrected *p*-values lower than 0.05 (*i.e*., FDR < 5%) were classified as *XP-EHH* outliers.

We additionally extracted information on the identity and functional annotation of all proteincoding genes annotated within these divergence/*XP-EHH* outlier windows (450 Kbp from the variant site). To further validate the identified candidate genes, we identified the location and ID of the nearest outlier *XP-EHH* site and calculated site-specific *EHH* in order to observe the span of *EHH* decay of that SNP in relation to the location of the gene. Gene ontology information for these genes was then categorized using the PANTHER database (Mi et al. 2021) and overlaps were identified between the candidates of selection identified via molecular evolution analysis.

## Data availability

The raw PacBio CLR and Hi-C Illumina reads used for the for *C. esox* and *C. gunnari* genome assemblies are available on NCBI under BioProject PRJNA857989. Per-sample, raw RADseq reads are available on NCBI BioProject PRJNA857737. RNAseq reads for *C. gunnari* and *C. esox* are available on NCBI BioProject PRJNA859929. For *C. esox, C. gunnari, and* the temperate outgroup *E. maclovinus*, the genome assemblies and associated annotations are hosted on <u>DRYAD</u>. Copies of the bioinformatic scripts used for analysis, including custom software, are available on <u>BitBucket</u>.

## Authors’ Contributions

AGR-C designed the experiments, collected specimens, constructed RADseq libraries, implemented bioinformatic data analyses, and wrote the manuscript. NR, BFM, and GM implemented bioinformatic data analyses. KTB and VY led RNAseq transcript sequencing. MH and SG collected and contributed specimens. C-HCC designed the experiments, collected specimens, led genome sequencing, analyzed the data, and edited the manuscript. JMC designed the experiments, implemented the bioinformatic data analyses, and edited the manuscript. All authors read and approved the manuscript.

## Acknowledgements

This work was supported by the National Science Foundation (NSF DGE 10-69157 IGERT to AGR-C, NSF OPP Grant 1645087 to JC and C-HCC, and NSF ANT Grant 11-42158 to C-HCC). The authors would like to thank Kira M. Long, Alida de Flamingh, and Carlos Ortiz-Alvarado for comments and discussions on the manuscript, Francesco Zapelloni and Jacob Anderson for assisting with the DNA extractions used for RAD sequencing, Shehani Gunawardena and Eliana Eng for their assistance in the bioinformatic analyses, Ernesto Davis Seguic for assistance on location in Chile, and Arthur DeVries for assistance with field collection and sampling.

## Notes

https://doi.org/10.5061/dryad.zgmsbccfd

### Competing Interest Statement

The authors have declared no competing interest.

### Summary of Updates

Manuscript revised following one initial round of review at Mol Biol & Evol. Aside from small editorial changes, the new version of the manuscript includes: * a map figure showing the species ranges * the number of all figures has changed due to the inclusion of the new map figure * editing of the results section to better describe the methods used (present later in the manuscript) * an inclusion of a gene family expansion analysis * an assessment of the presence/absence of circadian rhythm genes * re-writing of the discussion regarding circadian rhythm genes * clarifications on the version and reference datasets used for BUSCO analysis

