## Supplements for "Genomics of Secondarily Temperate Adaptation in the Only Non-Antarctic Icefish"

#### Methods

##### Optimization of contig-level genome assembly

For the assemblies of both *Champsocephalus esox* and *Champsocephalus gunnari*, we followed the assembly optimization methods described by (Rayamajhi et al. 2022), in which both genome assembly contiguity and gene-completeness are used as metrics of quality across different assembly iterations. Using our PacBio continuous long read (CLR) assemblies, we performed several contig-level assemblies iterating over different assemblers and input data parameters. First, we tested both the *FLYE* v2.5 (Kolmogorov et al. 2019) and *WTDBG2* v2.5 (Ruan and Li 2020) CLR assemblers. For each assembler, we tested different subsets of the input CLR data, subsampling reads across both different depths of coverage and length distributions (*i.e.*, only retaining within a specific size range). For each independent assembly iteration, we measured contiguity using *QUAST* v4.4 (Gurevich et al. 2013) and gene completeness using *BUSCO* v3.0.1 (Simão et al. 2015) with the `actinopterygii_odb9` gene dataset, selecting the initial assemblies yielding the best contiguity and completeness metrics.

In addition, we tested the effects of CLR read self-correction on the assemblies' gene-completeness. Previous results had indicated that an apparent excess in self-correction had detrimental effects, leading to an increase in the fragmentation and duplication of *BUSCO*s (*unpublished*). Therefore, to minimize these effects we assessed the gene-completeness of each assembly before any correction was applied, after single round of correction, and after two rounds of correction. Note that even when the *ARROW* correction is not performed, assemblies generated by *FLYE* will be internally self-corrected as part of the assembly pipeline.

For comparative purposes, using the optimization metrics described above, we retained two genome assemblies: the optimal assembly generated by *FLYE* and the optimal generated by *WTDGB2*. Based on contiguity and gene-completeness measures (Table S4), *FLYE* generated the best assemblies for both *C. esox* and *C. gunnari*. Hereby after, we refer to the *FLYE* assembly as the primary assembly for each species, while retaining the one generated by *WTDGB2* as secondary. These two assemblies, generated using independent assembly algorithms, allows us to perform manual curation of the assemblies at a later stage of the analysis. For example, structural variants were manually corroborated by confirming their presence in the two independent assemblies.

#### K-mer based annotation of AFGP sequences

To corroborate the manual annotation of antifreeze glycoprotein (AFGP) gene sequences in the reference assembly, *AFGP* sequences were additionally annotated using a k-mer based approach. First, we used an *AFGP* reference sequence, AFGP-H1A2 from NCBI (accession HQ447059.1), generated as part of the original characterization of the *AFGP* locus by (Nicodemus-Johnson et al. 2011), as a sequence query. We generated k-mers (k=17) for exons 1 and 2 of this reference sequence, which were then matched against the k-mers of a subject sequence of interest (*e.g.*, the reference assembly or the raw PacBio CLR reads). Matches between subject and query sequences were defined as k-mer "clumps", *i.e.*, regions with 20 or more individual matching k-mers located within 1 Kbp of one another. For either subject sequence, we annotated the coordinates of the corresponding *AFGP* k-mer clumps.

When using the reference assembly as the subject sequence, we compared the coordinates of the k-mer clumps against the location of the manually annotated sequences, an analysis comparable to the matching performed by BLAST. We additionally looked for k-mer clumps in the raw CLR reads, which we aligned to the reference assembly using *MINIMAP2* v2.24-r1122 (Li 2018). Using the coordinates from the alignments, we then compared the location the raw read *AFGP* k-mer clumps against the clumps located in the reference assembly. From the location of the alignments and clumps we confirmed that annotated *AFGPs* in the genome had corresponding *AFGP* k-mer clumps in the aligned reads and, *vice versa*, that *AFGP* k-mer clumps in the aligned raw reads had annotated *AFGP* sequences in the reference. We additionally visualized the alignments and clump location to confirm that the corresponding AFGP locus was span by the alignment of consecutive reads (*i.e.*, tiling). We only visualized alignments for reads larger than 15 Kbp in length, for which 75% or more of the total read length was aligned, and removing both non-primary and supplementary alignments. From these visualizations, we confirmed that there were no major patterns of disrupted alignments that might indicate the presence of assembly or scaffolding errors (*e.g.*, patterns of increase coverage due to collapsed repeats), verifying that regions with a reduction in read alignments corresponded to scaffolding gaps in the reference sequence. When validating the presence of both AFGP loci, in addition to the tiling, we inspected the presence of secondary/supplementary alignments to ensure that the raw reads uniquely mapped to their AFGP locus of origin.

### Figures

#### Figure S1: Conservation in genome structure and genome-wide synteny between *Champsocephalus* and the blackfin icefish *Chaenocephalus aceratus*

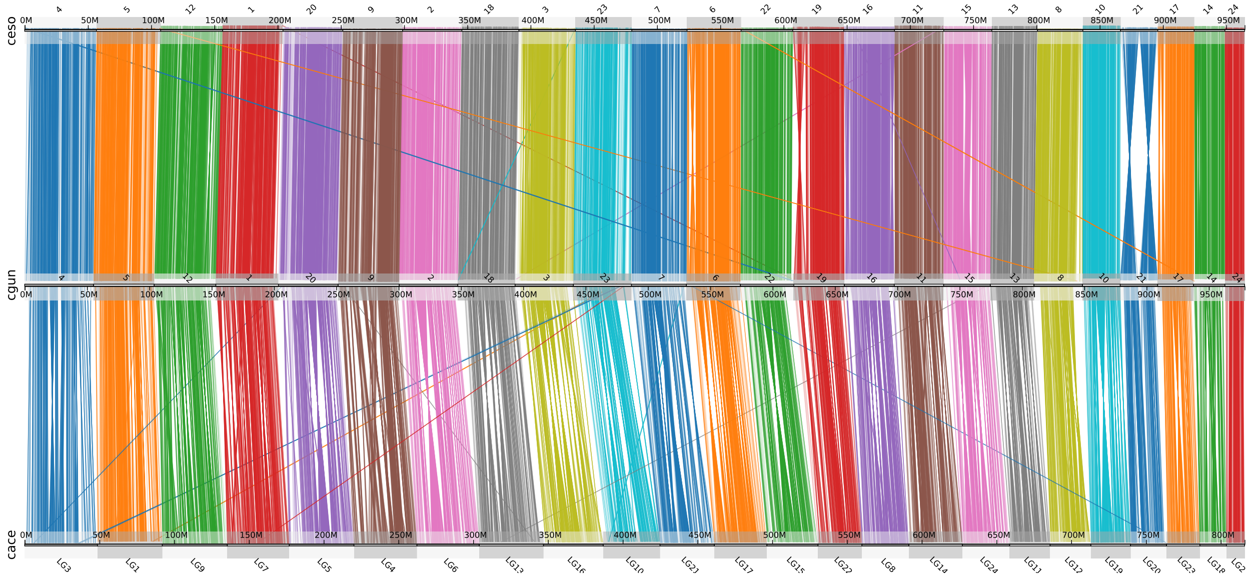


Genome-wide conserved synteny plots shows one-to-one correspondence between the 24 icefish chromosomes, reflecting a large-scale conservation in the organization of the genome. Top, middle, and bottom tracks correspond to the genome assemblies of *C. esox* (ceso), *C. gunnari* (cgun), and *C. aceratus* (cace) (Kim et al. 2019), respectively. Each line represents an orthologous gene between the two genomes, color-coded according to their chromosome of origin.

### Figure S2: Conservation in genome structure and genome-wide synteny between *Champsocephalus* and the South Georgia icefish *Pseudochaenichthys georgianus*

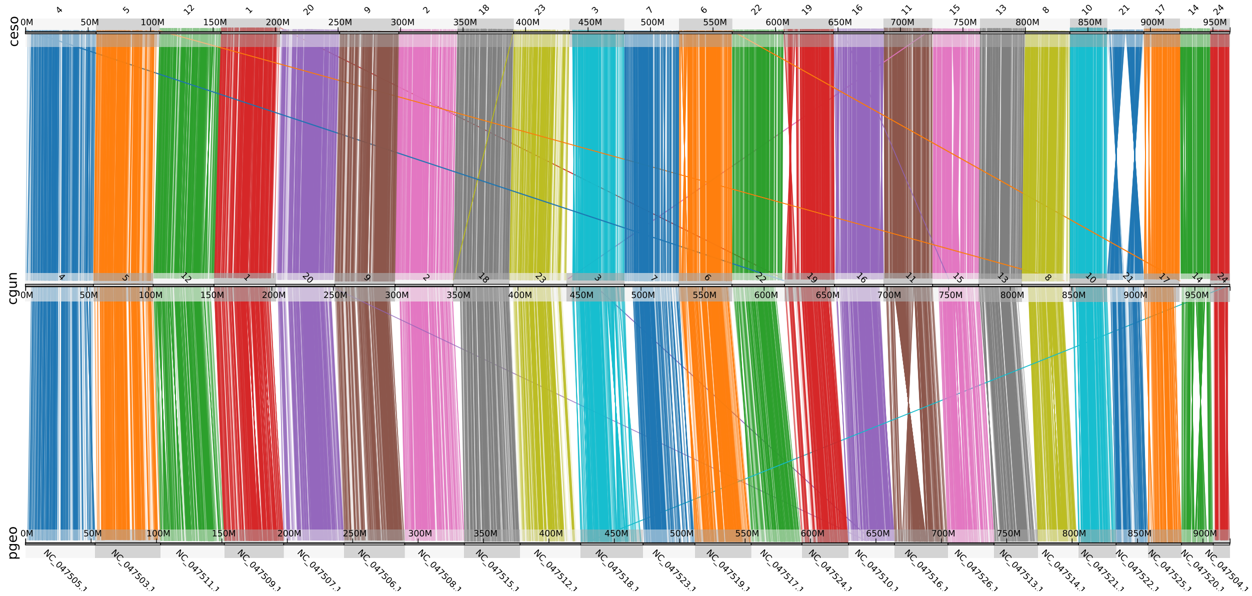


Genome-wide conserved synteny plots shows one-to-one correspondence between the 24 icefish chromosomes, reflecting a large-scale conservation in the organization of the genome. Top, middle, and bottom tracks correspond to the genome assemblies of *C. esox* (ceso), *C. gunnari* (cgun), and *P. georgianus* (pgeo) (Bista et al. 2022). Each line represents an orthologous gene between the two genomes, color-coded according to their chromosome of origin.

### Figure S3: Genetic divergence, genetic diversity, and selection across the genome

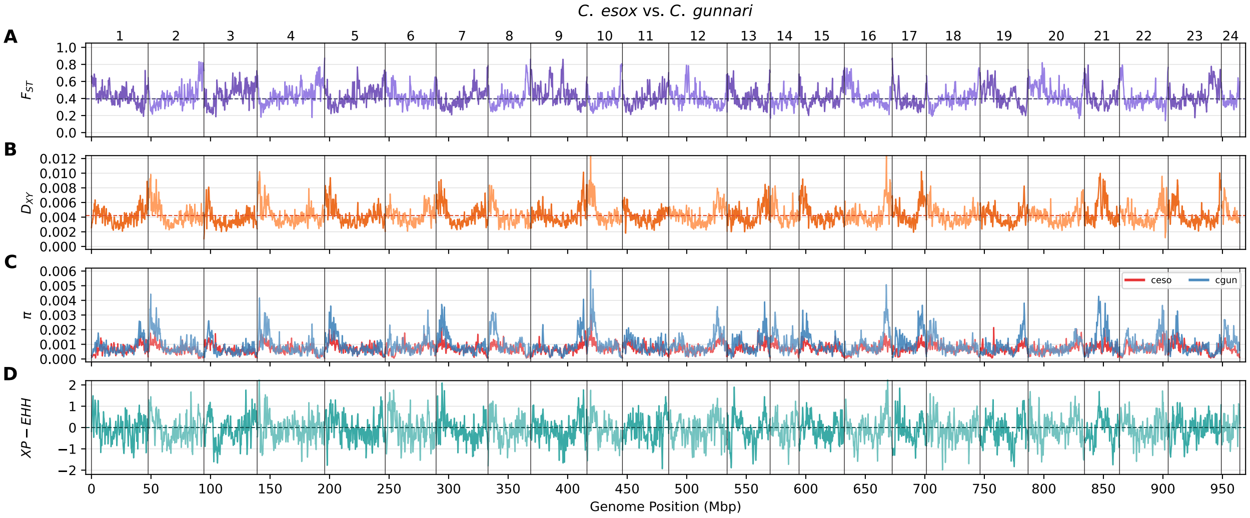


A) Relative genomic divergence ($F_{ST}$) between *C. esox* and *C. gunnari* populations. The X-axis shows the position along the *C. esox* reference genome. The boundaries between the chromosomes, which are sorted numerically, are shown by the vertical grey bars. The purple bar the kernel-smoothed $F_{ST}$ averaged across 150 Kbp windows. Dashed horizontal line shows the genome-wide average $F_{ST}$. B) Absolute genomic divergence ($D_{XY}$) between *C. esox* and *C. gunnari* populations. Orange line shows the kernel-smoothed average $D_{XY}$. Dashed horizontal line shows the genome-wide average $D_{XY}$. C) Nucleotide diversity ($\pi$) within the two studied icefish populations. The red and blue lines show the kernel-smoothed averages for *C. esox* and *C. gunnari*, respectively. D) Cross population extended haplotype homozygosity (*XP-EHH*) between *C. esox* and *C. gunnari* populations. Cyan lines shows the kernel-smoothed *XP-EHH* averaged over 150 Kbp windows. Positive and negative values reflect an increase in *EHH* in *C. esox* and *C. gunnari* populations, respectively.

#### Figure S4: Taxonomical sampling for the analysis of molecular evolution

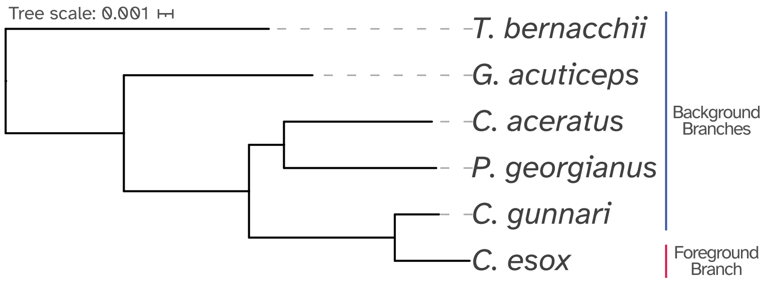


For the analysis of molecular evolution, we compared the orthologous coding sequences across six cryonotothenioid species, two red blooded Antarctic notothenioids (*Trematomus bernacchii* and *Gymnodraco acuticeps*), three Antarctic icefish (*C. gunnari*, *C. aceratus*, and *P. georgianus*), and our focal species *C. esox*. The phylogenetic relationship among these species, including both phylogenetic placement and branch distances, is based on the Notothenioidei-wide calibrated phylogeny generated by (Near et al. 2018). During the Branch-Site analysis of molecular evolution, *C. esox* was designated as the foreground branch to test for selection, while the five remaining species were labelled as background branches.

### Figure S5: Candidate genes adjacent to Extended Haplotype Homozygosity outlier SNPs

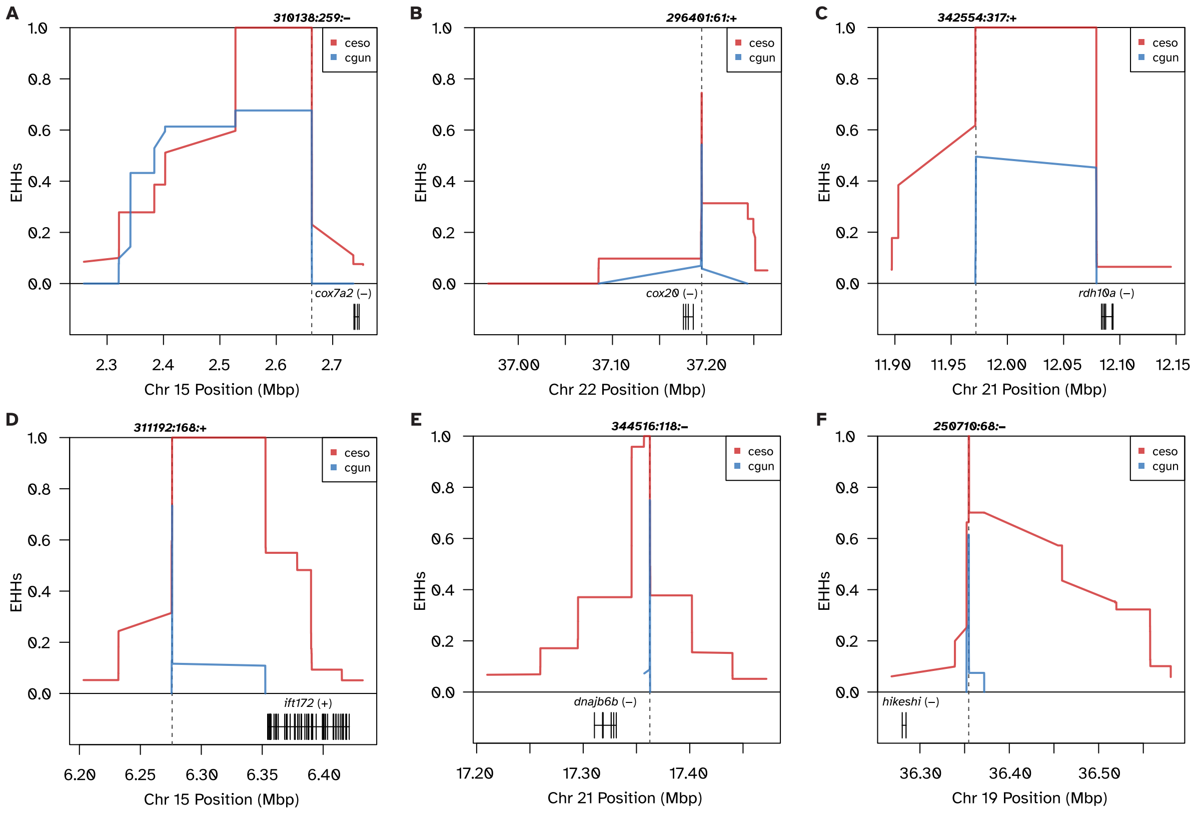


A) Signatures of population-level directional selection adjacent to the gene *cox7a2* (cytochrome c oxidase subunit 7A2b). The X axis shows the position on *C. esox* chromosome 15 in Mbp. The Y axis shows the site-specific Extended Haplotype Homozygosity (*EHHs*) values. The position of the target outlier SNP, labelled "310138:259:-" according to the *Stacks* catalog ID, is shown by the horizontal dashed line. The red and blue lines show the *EHHs* span for *C. esox and C. gunnari* populations, respectively. The location and stand of *cox7a2* is shown at the bottom. We see an increase in *EEHs* for *C. esox* populations downstream of *cox7a2*. B) *EHHs* adjacent to *cox20* (cytochrome c oxidase assembly factor COX20). Plot follows the annotation and color coding as described for panel A. We observe an increase in *EHHs* for the focal SNP in C. esox upstream to and spanning *cox20*. C) *EHHs* adjacent to *rdh10a* (retinol dehydrogenase 10a). Plot follows the annotation and color coding as described for panel A. An increase in *EEHs* or *C. esox* is observed upstream of *rdh10a*. D) *EEHs* adjacent to *ift172* (intraflagellar transport 172). Plot follows the annotation and color coding as described for panel A. We observe an increase in *EHHs* upstream of *ift172* in *C. esox*. E) *EHHs* adjacent to *dnajb6b* DnaJ heat shock protein family Hsp40 member B6b). Plot follows the annotation and color coding as described for panel A. We observe an increase in *EHHs* upstream of *ift172* in *C. esox* populations. F) *EHHs* adjacent to *hikeshi* (heat shock protein nuclear import factor hikeshi). Plot follows the annotation and color coding as described for panel A. We observe an increase in *EHHs* upstream of *hikeshi* in *C. esox* populations.

### Figure S6: Gene family expansion and contraction across notothenioids

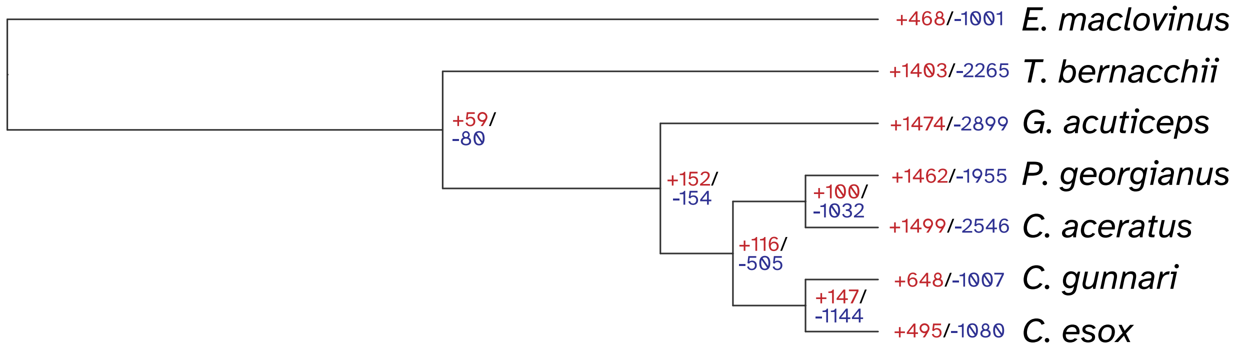


Ultrametric phylogenetic tree shows the relationship among seven notothenioid species, including a temperate outgroup (*Eleginops maclovinus*), two red blooded cryonotothenioids (*T. bernacchii* and *G. acuticeps*) and four icefishes (*P. georgianus*, *C. aceratus*, *C. gunnari*, and *C. esox*). For each node of the tree, the numbers indicate the number of significant gene family expansions (+, in red) and contractions (-, in blue).

### Figure S7: Effects of repeat annotation pipelines on repeat distributions

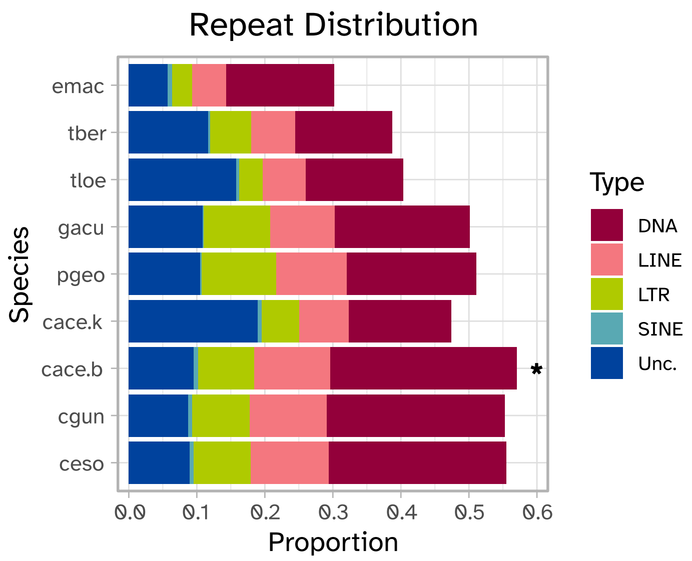


Interspersed repeat distributions for *C. esox* (ceso), *C. gunnari* (cgun), *C. aceratus* (cace), *P. georgianus* (pgeo), *G. acuticeps* (gacu), *T. bernacchii* (tber), *Trematomus loennbergii* (tloe), and *E. maclovinus* (emac). The colors in the bars represent proportions across five different repeat classes (Unc. = Unclassified). We show two repeat distributions for *C. aceratus*, cace.k which denotes the repeat distribution as published by (Kim et al. 2019), and cace.b (asterisk), a new repeat distribution generated for this study using the same *de novo* repeat library and *REPEATMASKER/REPEATMODELER* annotation pipeline implemented for *C. esox* and *C. gunnari*. The updated annotation pipeline results in a total increase in identified repeats (similar to the distribution observed in *C. esox* and *C. gunnari*) and a reduction in the frequency of unclassified elements. Overall, this suggest that the higher repeat content observed for *C. esox* and *C. gunnari* are not unique to members of the genus *Champsocephalus* and might be instead product of the repeat annotation pipeline utilized.

### Figure S8: AFGP loci and genomic rearrangements in chromosome 3

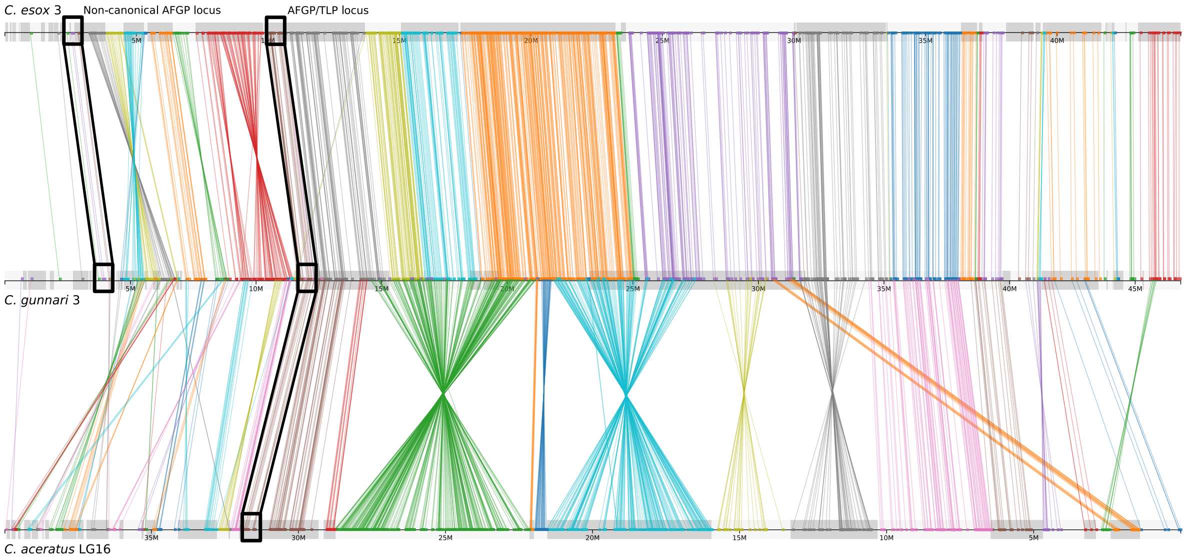


Conserved synteny plot showing patterns of orthologous gene clusters. Top, middle, and bottom tracks correspond to *C. esox* chromosome 3, *C. gunnari* chromosome 3, and *C. aceratus* chromosome L16, respectively. Each line represents an orthologous gene between the genomes, color-coded based on their contig location on the corresponding query chromosome (*C. esox* 3 in the top comparison, *C. aceratus* LG16 on the bottom). Black boxes denote the approximate locations of the two AFGP loci, the non-canonical locus (present only in *C. esox* and *C. gunnari*) and the canonical AFGP/TLP locus present in the three species.

#### Figure S9: Loss of circadian-specific genes across notothenioid genomes

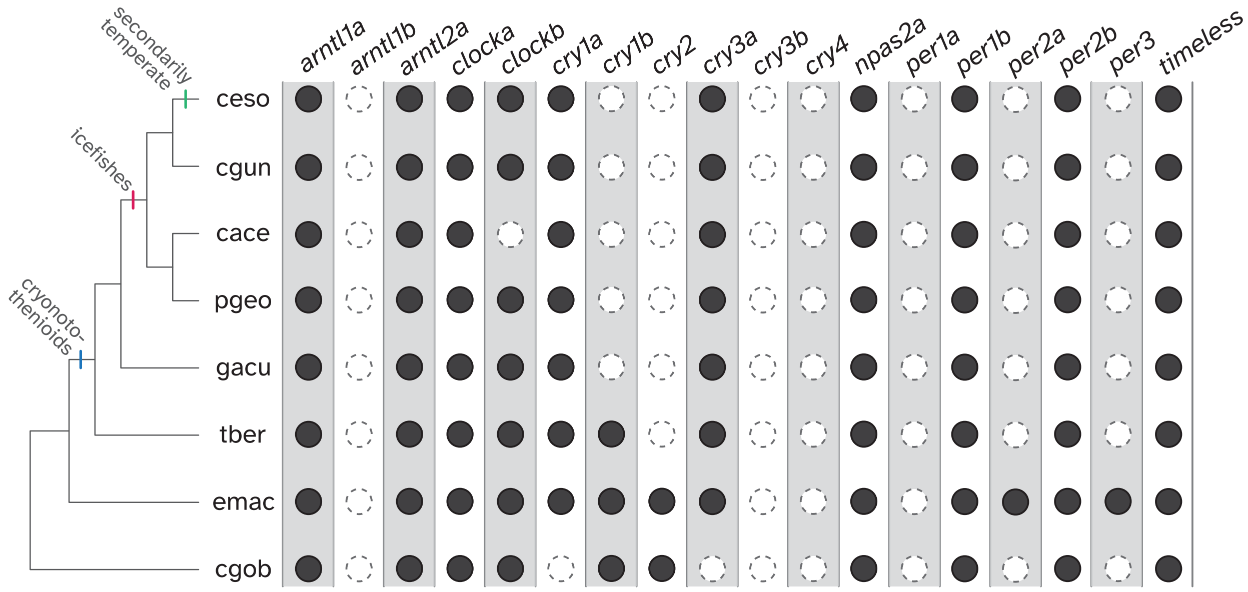


Presence or absence of circadian genes across eight notothenioid species, *C. esox* (ceso), *C. gunnari* (cgun), *C. aceratus* (cace), *P. georgianus* (pgeo), *G. acuticeps* (gacu), *T. bernacchii* (tber), and *E. maclovinus* (emac), *Cottoperca gobio* (cgob). Phylogenetic relationship among the species is shown in the tree on the left. Solid black circles and empty white circles show presence and absence, respectively, of the corresponding ortholog in the annotated notothenioid genome. The nomenclature of circadian gene orthologs follows the designation used by (Toloza-Villalobos et al. 2015; Kim et al. 2019). For a given gene, the orthology between the reference and query notothenioid species was determined by reciprocal *BLAST* best hit, as specified by SYNOLOG (Catchen et al. 2009). See Table S9 for the specification of the reference circadian gene sequences and orthologs coordinates in the queried notothenioid genomes.

### Figure S10: Molecular evolution in GNAT2

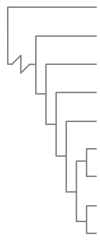

```
drer_gnat2   YTKEEQMEFRSIIFGNILQSALAIIRGMEMLSINFGSPSAQEDSQKLQNLSDSIEEGTMP   120
cgob_gnat2   YSRQEQLEFRGVVYGNILQSALAIIRGMEMLSIDFGSTAAQENAQKLQNLSDSIEEGTMP   116
emac_gnat2   YTKEEQLEFRAVIYGNILQSALAIIRGMEMLTIDFGSSSSQEDAQKLQNLSDSIEEGTMP   116
tber_gnat2   YSKEEQLEFRGVVYGNILQSALAIIRGMEMLTIDFGSAAAQENAQKLQNLSDSIEEGTMP    79
gyac_gnat2   YTKEEQLEFRGVVYGNILQSTLAIIRGMEMLTIDFGSPTAQENAQKLQNLSDSIEEGTMP   116
cace_gnat2   YTKEEQLEFRGIVFGNILQSTLAIIRGMEMLNIDFGSPTAQENAQKLQNLSDSIEEGTMP   116
pgeo_gnat2   YTKEEQLEFRGIVFGNILQSTLAIIRGMEMLTIDFGSPTAQENAQKLQNLSDSIEEGTMP   116
cgun_gnat2   YTKEEQLEFRGIVFGNILQSALAIIRGMEMLTIDFGSPAAQENAQKLQNLSDSIEEGTMP   116
ceso_gnat2   YTKEEQLEFRTIVFGNILQSALAIIRGMEMLTIDFGSPAAQENAQKLQNLSDSIEEGTMP   116
             *:::**:***  :::******:**********.*:***  ::**::****************
```

Alignment for the coding sequence of the gene *gnat2* (G protein subunit alpha transducin 2) between the zebrafish *Danio rerio* (drer), *C. gunnari* (cgun), *C. esox* (ceso), *C. aceratus* (cace), *P. georgianus* (pgeo), *T. bernacchii* (tber), *G. acuticeps* (gyac), *E. maclovinus* (emac), and *C. gobio* (cgob). Phylogenetic relationship is shown by the cladogram on the left. The figure only shows a partial alignment of 60-amino acids (end position shown by the numbers on the right). The purple row highlights the sequence for *C. esox,* while the cyan column highlights the candidate site under selection in this species. In contrast to all other notothenioids assessed, which contain a Gly (G) or Ala (A) residue at the selected site, *C. esox* shows a Thr (T).

### Figure S11: Characterization and curation of the non-canonical AFGP locus in *C. esox*

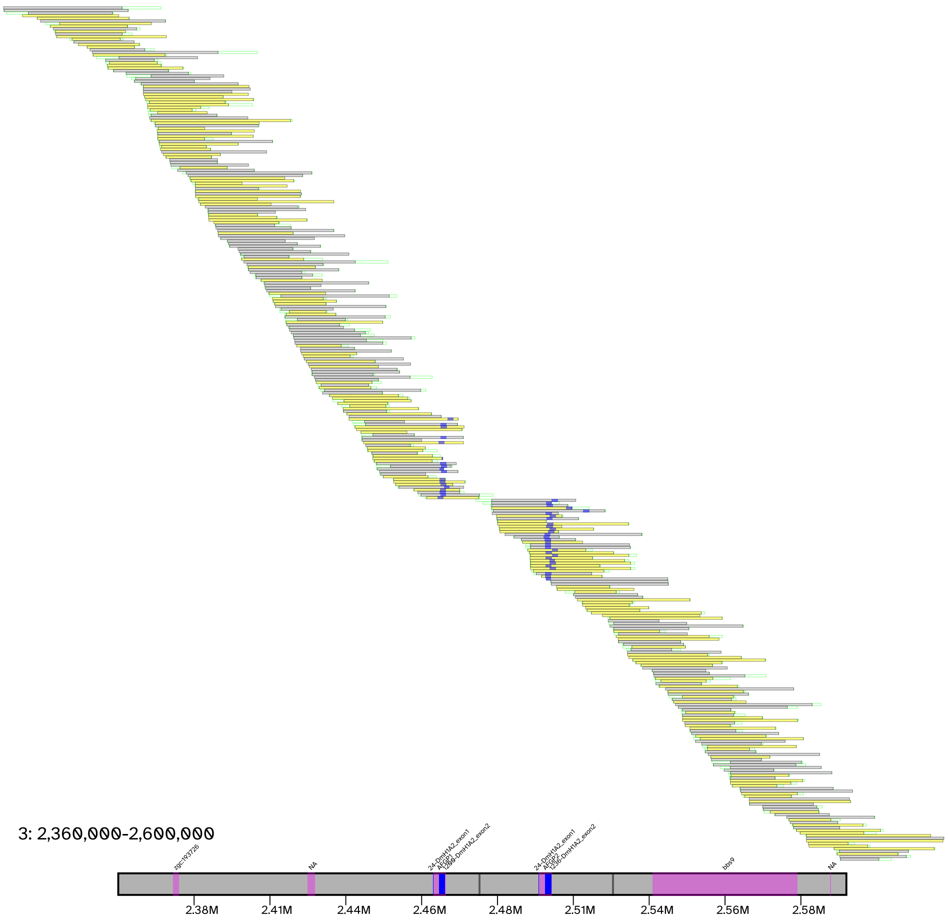


Tiling of raw reads and location of *AFGP* k-mer clumps in the non-canonical AFGP locus in *C. esox*. The small horizontal bars spanning the image represent the aligned reads. Grey lines refer to reads mapping in the forward orientation, while yellow lines refer to reads mapping on the reverse orientation. Green boxes denote portions of the reads soft-clipped during alignment. Blue boxes show the approximate location of the *AFGP* clumps. The bottom horizontal bar represents the underlying chromosome. Pink boxes denote the annotated protein-coding genes in this region. As in the raw reads, blue boxes denote the location of the *AFGP* clumps. Vertical dark grey bars represent gaps in the reference sequence. We display alignments for the region between 2.36 to 2.60 Mbp on chromosome 3. At both ends of this region we see *zgc:193726* and *bbs9*, the two annotated genes that form the 5' and 3' boundaries of the locus and display conserved synteny to *C. gunnari*. Two *AFGP* copies are seen in blue. There is one scaffolding gap located around 2.475 Mbp, located between the two *AFGP* copies.

### Figure S12: Characterization and curation of the canonical AFGP/TLP locus in *C. esox*

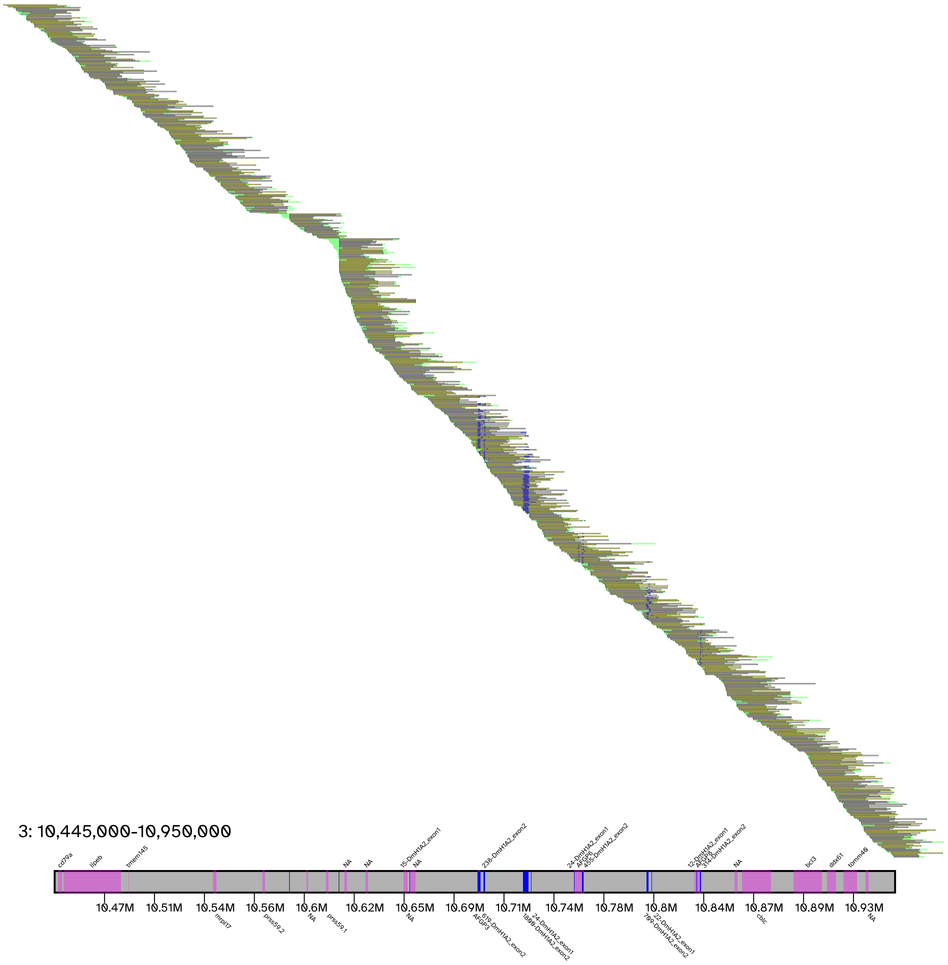


Tiling of raw reads and location of *AFGP* k-mer clumps in the canonical AFGP/TLP locus in *C. esox*, displaying alignments in between 10.445 and 10.95 Mbp on chromosome 3. Figure follows annotation and color coding as described in Fig. S7. At both ends of the region we see *hsl* (*lipeb* homolog) and *tomm40*, which define the 5' and 3' end boundaries of the canonical locus as described by (Nicodemus-Johnson et al. 2011). We observe a total of 7 *AFGP* clumps. The first one at around 10.65 Mbp is only composed of AFGP exon 1 and represents a copy of *TLP.* The next two clumps, at around 10.7 Mbp are only composed of AFGP exon 2 and represent the two partial *AFGP* copies. The four remaining clumps contain both AFGP exon 1 and exon 2 k-mers, and represent the complete, albeit pseudogenized, *AFGP* copies. There are two scaffolding gaps, one at around 10.58 Mbp and another at 10.61 Mbp, which denote a small contig containing two annotated *tryp1* copies.

### Figure S13: Characterization and curation of the non-canonical AFGP locus in *C. gunnari*

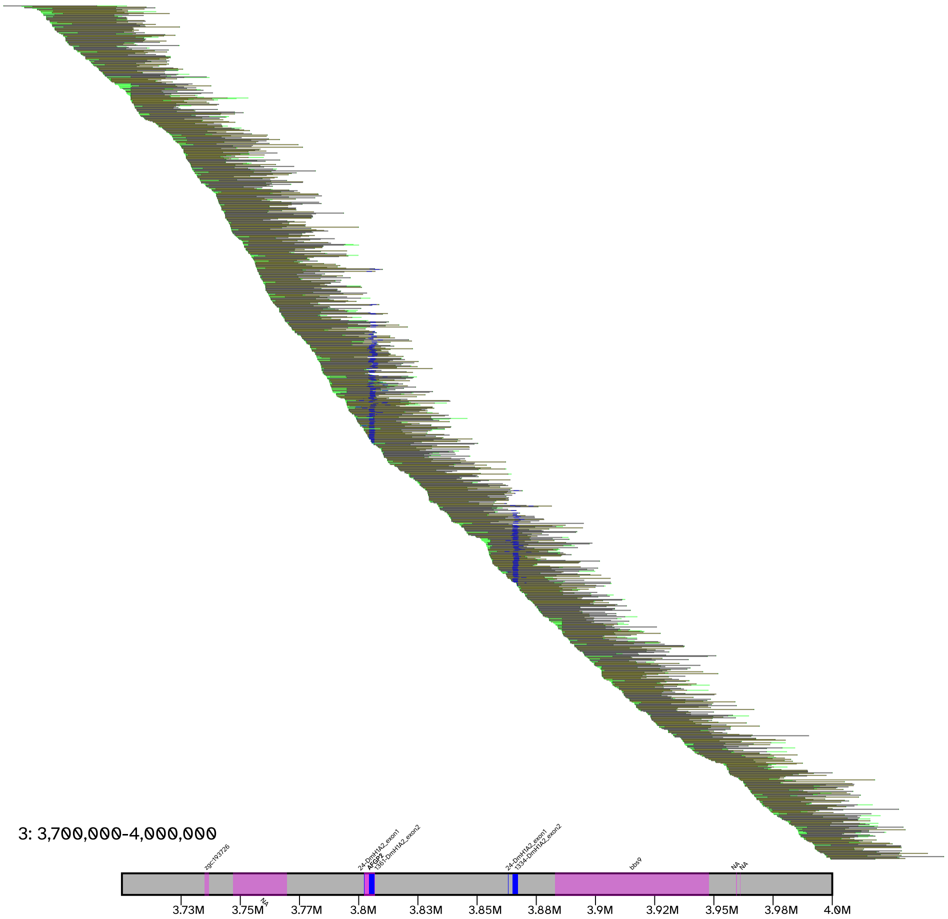


Tiling of raw reads and location of *AFGP* k-mer clumps in the non-canonical AFGP locus in *C. gunnari*, displaying alignments in between 3.7 to 4.0 Mbp on chromosome 3. The figure follows annotation and color coding as described in Fig. S7. At both ends of this genomic region we see *zgc:193726* and *bbs9*, the two annotated genes that form the 5' and 3' boundaries of the locus and display conserved synteny to *C. gunnari*. Two *AFGP* copies are seen in blue.

### Figure S14: Characterization and curation of the canonical AFGP/TLP locus in *C. gunnari*

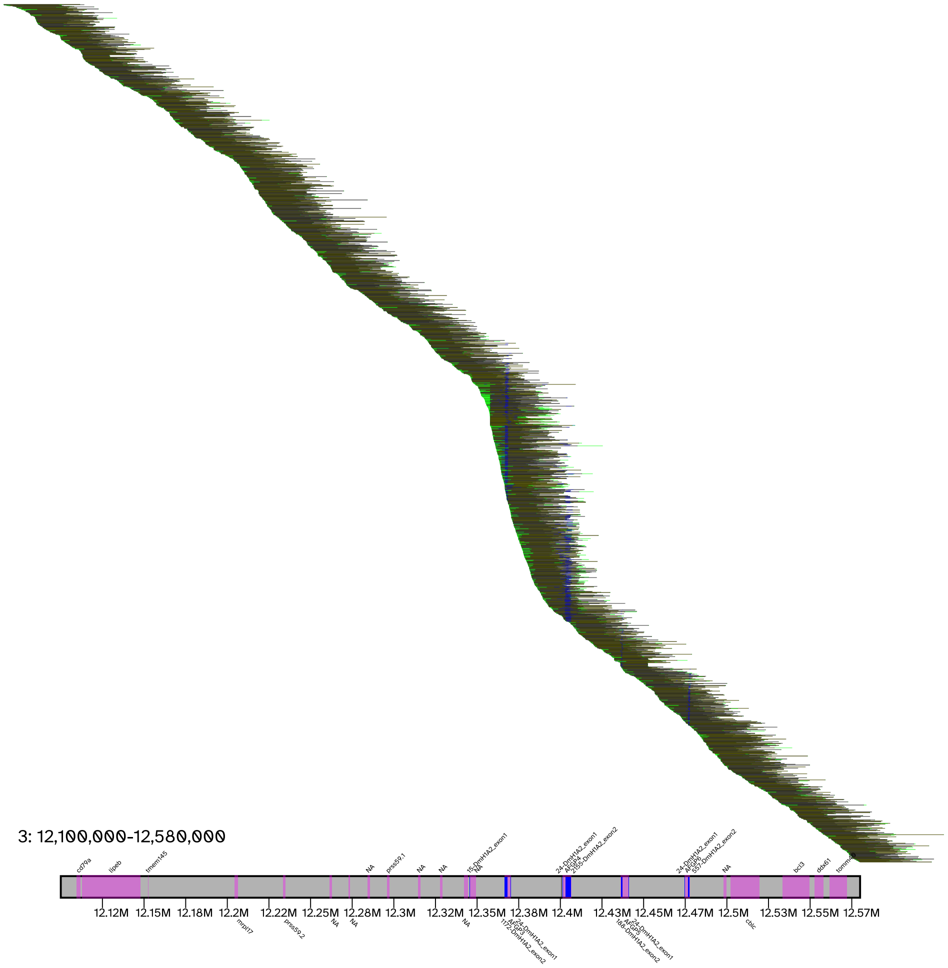


Read tiling of the canonical AFGP/TLP locus in *C. gunnari*, displaying alignments in between 12.1 and 10.580 Mbp on chromosome 3. Figure follows annotation and color coding as described in Fig. S7. At both ends of the region we see *hsl* (*lipeb* homolog) and *tomm40*, which define the canonical 5' and 3' end boundaries locus (Nicodemus-Johnson et al. 2011). We observe a total of 5 major AFGP clumps. The first one at around 12.34 Mbp is only composed of *AFGP* exon 1 and represents a copy of *TLP.* The four remaining clumps contain both *AFGP* exon 1 and exon 2 k-mers and represent complete *AFGP* copies. There is one scaffolding gap at around 12.4 Mbp, but no disruption in the tiling of reads is observed.

### Tables

#### Table S1: Gene completeness assessment of the chromosome-scale genome assemblies

| Species | *C. esox* | *C. gunnari* |
|---|---|---|
| ***BUSCO*** **v3.0.1 (genome mode)** | | |
| Complete | 4,400 (96.0%) | 4,404 (96.1%) |
| Complete & Single Copy | 4,298 (93.8%) | 4,296 (93.7%) |
| Complete & Duplicated | 102 (2.2%) | 108 (2.4%) |
| Fragmented | 52 (1.1%) | 56 (1.2%) |
| Missing | 132 (2.9%) | 124 (2.7%) |
| Total | 4,584 | 4,584 |
| Reference Gene Set | `actinopterygii_odb9` | `actinopterygii_odb9` |
| ***BUSCO*** **v5.1.3 (genome mode)** | | |
| Complete | 3,506 (96.3%) | 3,506 (96.3%) |
| Complete & Single Copy | 3,463 (95.1%) | 3,469 (95.3%) |
| Complete & Duplicated | 43 (1.2%) | 37 (1.0%) |
| Fragmented | 14 (0.4%) | 19 (0.5%) |
| Missing | 120 (3.3%) | 115 (3.2%) |
| Total | 3,640 | 3,640 |
| Reference Gene Set | `actinopterygii_odb10` | `actinopterygii_odb10` |
| ***BUSCO*** **v5.1.3 (protein mode)** | | |
| Complete | 3,319 (91.2%) | 3,358 (92.3%) |
| Complete & Single Copy | 3,120 (85.7%) | 3,187 (87.6%) |
| Complete & Duplicated | 199 (5.5%) | 171 (4.7%) |
| Fragmented | 132 (3.6%) | 125 (3.4%) |
| Missing | 189 (5.2%) | 157 (4.3%) |
| Total | 3,640 | 3,640 |
| Reference Gene Set | `actinopterygii_odb10` | `actinopterygii_odb10` |

### Table S2: Assembly statistics of notothenioid genomes used in the study

| Species | *C. esox* | *C. gunnari* | *C. aceratus* | *P. georgianus* | *G. acuticeps* | *T. bernacchii* | *T. loennbergii* | *E. maclovinus* |
|---|---|---|---|---|---|---|---|---|
| Total Size (bp) | 987,108,782 | 994,201,109 | 1,065,748,709 | 1,026,101,545 | 996,888,495 | 867,125,071 | 1,210,523,816 | 606,289,673 |
| Total Contig Size (bp) | 986,435,181 | 993,506,282 | 1,065,645,263 | 1,025,462,058 | 996,695,919 | 867,007,390 | 1,210,488,816 | 606,099,673 |
| Percentage Contigs | 99.932% | 99.930% | 99.990% | 99.938% | 99.981% | 99.986% | 99.997% | 99.969% |
| Number of Contigs | 3,499 | 2,992 | 3,853 | 4,021 | 4,183 | 1,793 | 3,919 | 406 |
| Number of Scaffolds | 2,067 | 1,536 | 2,821 | 1,563 | 2,618 | 864 | 3,569 | 26 |
| Largest Contig (bp) | 9,841,188 | 15,571,738 | 9,422,831 | 4,038,360 | 4,294,025 | 1,402,669 | 17,998,618 | 19,993,184 |
| Largest Scaffold (bp) | 56,719,351 | 55,323,500 | 48,701,853 | 53,425,585 | 13,140,105 | 8,748,152 | 48,466,630 | 37,057,500 |
| Contig N50 | 2,611,138 | 3,182,242 | 1,497,474 | 661,283 | 534,678 | 601,448 | 968,532 | 7,582,354 |
| Scaffold N50 | 43,613,322 | 44,089,792 | 34,368,030 | 42,832,159 | 1,921,211 | 2,904,264 | 15,527,446 | 26,674,500 |
| Contig L50 | 107 | 84 | 170 | 431 | 503 | 160 | 232 | 27 |
| Scaffold N50 | 11 | 11 | 14 | 11 | 132 | 27 | 22 | 11 |
| Number of Chromosome-Scale Scaffolds | 24 | 24 | 24 | 24 | -- | -- | -- | 24 |
| Total Bases in Chromosomes | 964,606,985 | 978,689,797 | 816,521,294 | 922,268,569 | -- | -- | -- | 606,192,173 |
| Percent Assembly in Chromosomes | 97.720% | 98.440% | 76.615% | 89.881% | -- | -- | -- | 99.984% |
| Study | This Study | This Study | Kim et al. 2019 | Bista et al. 2022 | Bista et al. 2022 | Bista et al. 2022 | Jo et al. 2021 | This study/Cheng et al. *in prep* |

#### Table S3: Distribution of repeats in the *C. esox* and *C. gunnari* genomes

| | ***C. esox*** | | ***C. gunnari*** | |
|---|---|---|---|---|
| | Total Bases (bp) | Percentage of assembly | Total Bases (bp) | Percentage of assembly |
| Total Repeats | 586,145,195 | 59.38% | 591,052,559 | 59.45% |
| LTR | 83,410,692 | 8.45% | 84,407,674 | 8.49% |
| SINE | 5,527,809 | 0.56% | 5,865,787 | 0.59% |
| LINE | 112,629,112 | 11.41% | 113,140,086 | 11.38% |
| DNA | 257,635,392 | 26.10% | 259,784,750 | 26.13% |
| Unclassified | 88,543,658 | 8.97% | 86,396,076 | 8.69% |

#### Table S4: Population-level measures of genetic diversity and divergence across two icefish species

| **Population A** | **Population B** | **PopA $\pi$** | **PopB $\pi$** | $F_{ST}$ | $D_{XY}$ |
|---|---|---|---|---|---|
| *C. gunnari* (South Georgia) | *C. gunnari* (West Antarctic Peninsula) | 0.12 | 0.1 | 0.092 | 0.0021 |
| *C. esox* (Puerto Natales) | *C. esox* (Canal Bárbara) | 0.078 | 0.077 | 0.055 | 0.0014 |
| *C. esox* (Puerto Natales + Canal Bárbara) | *C. gunnari* (South Georgia) | 0.088 | 0.12 | 0.4 | 0.0042 |

#### Table S5: *PAML* results for the *dN*/*dS* candidate genes under positive selection

[Supplementary Excel file]

#### Table S6: Divergence/*XP-EHH* windows candidate genes and location of adjacent outlier *XP-EHH* sites

[Supplementary Excel file]

#### Table S7: Functional annotation for all candidate genes according to their corresponding zebrafish ortholog

[Supplementary Excel file]

### Table S8: Top 30 candidates for gene family contraction and expansion in *C. esox*

| Family ID[1] | Count[2] | Change[3] | Status | *p*-value[4] | Description[5] |
|---|---|---|---|---|---|
| N0.HOG0000105 | 0 | -7 | contraction | 2.50E-15 | Zona Pellucida |
| N0.HOG0002556 | 7 | 6 | expansion | 1.80E-13 | -- |
| N0.HOG0000104 | 0 | -6 | contraction | 2.11E-13 | Reverse Transcriptase-associated |
| N0.HOG0000182 | 12 | 6 | expansion | 7.79E-11 | -- |
| N0.HOG0000635 | 7 | 5 | expansion | 1.27E-10 | -- |
| N0.HOG0000700 | 7 | 5 | expansion | 1.27E-10 | -- |
| N0.HOG0000884 | 7 | 5 | expansion | 1.27E-10 | Histone Methyltransferase |
| N0.HOG0001231 | 7 | 5 | expansion | 1.27E-10 | -- |
| N0.HOG0001420 | 7 | 5 | expansion | 1.27E-10 | -- |
| N0.HOG0001076 | 5 | 4 | expansion | 2.53E-09 | BED Zinc Finger |
| N0.HOG0000163 | 0 | -4 | contraction | 4.80E-09 | Myosin |
| N0.HOG0000750 | 8 | 4 | expansion | 9.02E-08 | -- |
| N0.HOG0001050 | 4 | 3 | expansion | 3.00E-07 | Myosin H |
| N0.HOG0001254 | 4 | 3 | expansion | 3.00E-07 | Lysozyme G |
| N0.HOG0002584 | 4 | 3 | expansion | 3.00E-07 | UNC-5 Netrin Receptor |
| N0.HOG0003545 | 4 | 3 | expansion | 3.00E-07 | Proteasome 26S |
| N0.HOG0004795 | 4 | 3 | expansion | 3.00E-07 | Formyl Peptide Receptor |
| N0.HOG0000685 | 0 | -3 | contraction | 3.74E-07 | Harbinger Transposase |
| N0.HOG0001366 | 0 | -3 | contraction | 3.74E-07 | -- |
| N0.HOG0004783 | 5 | 3 | expansion | 1.19E-06 | BED Zinc Finger |
| N0.HOG0000463 | 1 | -3 | contraction | 1.35E-06 | PR/SET Domain |
| N0.HOG0000301 | 2 | -3 | contraction | 3.26E-06 | Odorant Receptor, Family F |
| N0.HOG0000326 | 2 | -3 | contraction | 3.26E-06 | Microglobulin/Bikunin Precursor |
| N0.HOG0000099 | 3 | -3 | contraction | 6.37E-06 | Microtubule Binding Proteins |
| N0.HOG0000567 | 7 | 3 | expansion | 8.15E-06 | Myosin H |
| N0.HOG0000021 | 8 | 3 | expansion | 1.58E-05 | -- |
| N0.HOG0000016 | 3 | 2 | expansion | 3.56E-05 | -- |
| N0.HOG0000022 | 3 | 2 | expansion | 3.56E-05 | -- |
| N0.HOG0000094 | 3 | 2 | expansion | 3.56E-05 | -- |
| N0.HOG0000270 | 3 | 2 | expansion | 3.56E-05 | UNC-13 Homolog |

[1]N0 Phylogenetic Hierarchical Orthogroup ID from *ORTHOFINDER*; [2]Number of ortholog identified in *C. esox*; [3]Changes in ortholog number in *C. esox* in relation to ancestral node; [4]As calculated by *CAFÉ* v5; [5]According to the *INTERPROSCAN* functional annotation.

#### Table S9: Circadian rhythm gene orthologs in notothenioid genome annotations

[Supplementary Excel file]

#### Table S10: Meta data of specimens used for sequencing

| | ***C. gunnari*** | ***C. esox*** |
|---|---|---|
| **Genome/Hi-C sequencing (1 individual)** | | |
| Catch site | Gerlache Strait, Antarctica | Puerto Natales, Chile |
| GPS | -63.45, -62.50 | -51.73, -72.52 |
| Catch date | July, 2014 | December, 2017 |
| Specimen id | UIUC-203 | UIUC-362 |
| Weight | 367 g | 93 g |
| Total Length | 38.8 cm | 27.9 cm |
| Sex | male | male |
| Tissue for DNA | white muscle | isolated hepatocytes |
| **RAD sequencing** | | |
| Catch sites and dates | (i) South Georgia, Antarctica (February 2017) | (i) Puerto Natales, Chile (January 2008; October 2009; December 2017) |
| | (ii) Low Island, Dallman Bay, Gerlache Strait, Antarctica (June-July, 2008, 2014) | (ii) Canal Bárbara, Chile (April 2016) |
| **Transcript sequencing** | | |
| Catch sites and dates | Low Island, Dallman Bay, Gerlache Strait, Antarctica (June-July, 2008, 2014)[1] | Puerto Natales, Chile (June 2008; October 2009)[1] |

[1]Subset of the same individuals used for RAD sequencing

#### Table S11: PacBio CLR genome sequencing statistics - 2 SMRT cells per species

| | ***C. esox*** | ***C. gunnari*** |
|---|---|---|
| Total number of reads | 10,734,722 | 12,602,890 |
| Total read yield (Mbp) | 141,014.63 | 17,073.81 |
| Mean read length (bp) | 13,136.31 | 29,775.00 |
| Subread N50 (bp) | 24,348 | 29,775 |
| Number of reads >20Kbp | 2,339,419 | 4,688,701 |
| Number of reads >50Kbp | 301,544 | 159,328 |

Table S12: RQN (RNA quality number) of tissue RNA samples used in paired-end (2x250bp) transcript sequencing on Illumina HiSeq2500

| ***C. esox*** | | ***C. gunnari***[1] | |
|---|---|---|---|
| **Tissue** | **RQN** | **Tissue** | **RQN** |
| Brain | 8.9 | Brain | 8.4 |
| Eye cup | 7.4 | Eye cup | 7.9 |
| Lens | 8.6 | Eye lens | 9.4 |
| Gill | 7.7 | Gills | 9.5 |
| Heart (ventricle) | 7.5 | Heart | 9.2 |
| Liver | 9 | Liver | 9.2 |
| Anterior stomach | 6.6 | Esophagus stomach | 9.3 |
| -- | -- | Pyloric ceca | 9.9 |
| -- | -- | Mesentery | 9.8 |
| -- | -- | Anterior intestine | 9.8 |
| Head kidney | 6.5 | Head kidney | 9.1 |
| -- | -- | Red body | 8.9 |
| Renal kidney | 7.4 | Caudal kidney | 9.1 |
| Spleen | 8 | Spleen | 9.5 |
| Red muscle | 7.1 | Red muscle | 9.0 |
| White muscle | 8.8 | White muscle | 9.5 |
| Skin | 7.7 | Skin | 9.2 |
| Testis | 9.4 | Testis | 10.0 |
| -- | | Ovary | 9.8 |
| **Total pool[2]** | **7.7** | **Total pool[2]** | **9.2** |

[1]*C. gunnari* individuals sampled from the West Antarctic Peninsula (see Table S10). [2]Pool composed of 1 µg RNA from each tissue.

### Table S13: Statistics of contig-level genome assemblies

| Species | ***C. esox*** | | ***C. gunnari*** | |
|---|---|---|---|---|
| Assembler | *FLYE* | *WTDBG2* | *FLYE* | *WTDBG2* |
| Read depth (x) | 70 | 80 | 80 | 80 |
| Min. read length (Kbp) | 10 | 15 | 15 | 15 |
| Max. read length (Kbp) | 40 | 40 | -- | 40 |
| Assembly size (Gbp) | 0.986 | 1.04 | 0.993 | 1.04 |
| N50 (Mbp) | 3.1 | 1.9 | 3.7 | 3.2 |
| Number of fragments | 3,210 | 5,680 | 2,777 | 5,061 |
| Largest fragment (Mbp) | 15.3 | 16.2 | 21 | 22.4 |
| *BUSCO*[1] v3.0.1 Complete | 96.30% | 95.70% | 96.50% | 95.80% |
| *BUSCO*[1] v3.0.1 Single | 93.80% | 93.50% | 93.70% | 93.60% |
| *BUSCO*[1] v3.0.1 Duplicated | 2.50% | 2.20% | 2.80% | 2.20% |
| *BUSCO*[1] v3.0.1 Fragmented | 1.10% | 1.10% | 1.00% | 1.00% |
| *BUSCO** v3.0.1 Missing | 2.60% | 3.20% | 2.50% | 3.20% |
| Rounds of *ARROW* correction | 0 | 1 | 0 | 1 |
| Status | Primary | Secondary | Primary | Secondary |

[1]*BUSCO* analysis with the `actinopterygii_odb9` gene dataset
